# Annotating publicly-available samples and studies using interpretable modeling of unstructured metadata

**DOI:** 10.1101/2024.06.03.597206

**Authors:** Hao Yuan, Parker Hicks, Mansooreh Ahmadian, Kayla Johnson, Lydia Valtadoros, Arjun Krishnan

**Affiliations:** Michigan State University, Genetics and Genome Sciences Program, East Lansing, 48823, USA; Michigan State University, Ecology, Evolution, and Behavior Program, East Lansing, 48823, USA; University of Colorado Anschutz Medical Campus, Department of Biomedical Informatics, Aurora, 80045, USA; University of Colorado Anschutz Medical Campus, Department of Biostatistics and Informatics, Aurora, 80045, USA

## Abstract

Reusing massive collections of publicly available biomedical data can significantly impact knowledge discovery. However, these public samples and studies are typically described using unstructured plain text, hindering the findability and further reuse of the data. To combat this problem, we propose *txt2onto 2*.*0*, a general-purpose method based on natural language processing and machine learning for annotating biomedical unstructured metadata to controlled vocabularies of diseases and tissues. Compared to the previous version (*txt2onto 1*.*0*), which uses numerical embeddings as features, this new version uses words as features, resulting in improved interpretability and performance, especially when few positive training instances are available. *Txt2onto 2*.*0* uses embeddings from a large language model during prediction to deal with unseen-yet-relevant words related to each disease and tissue term being predicted from the input text, thereby explaining the basis of every annotation. We demonstrate the generalizability of *txt2onto 2*.*0* by accurately predicting disease annotations for studies from independent datasets, using proteomics and clinical trials as examples. Overall, our approach can annotate biomedical text regardless of experimental types or sources. Code, data, and trained models are available at https://github.com/krishnanlab/txt2onto2.0.

**Key points:**

- We developed *txt2onto 2*.*0*, a computational method that combines language models and machine learning to annotate public samples and studies with standardized tissue and disease terms, with a focus on interpretability and explainability.
- *Txt2onto 2*.*0* uses word/phrase occurrence statistics to represent sample/study metadata, train machine learning models, and predict terms in controlled vocabularies to annotate each sample and study. This approach allows the model to keep track of predictive words related to model decisions and easily separate informative from uninformative words.
- *Txt2onto 2*.*0* outperforms its predecessor, *txt2onto 1*.*0*, in tissue and disease annotation, especially when training data is limited.
- The predictive features learned by *txt2onto 2*.*0* are highly interpretable. These features not only include explicit mentions of the actual disease or tissue terms but also related biomedical concepts, including words that are unseen by the model during training.
- Although trained on metadata of transcriptomes, *txt2onto 2*.*0* is capable of annotating disease and tissue for any kind of biomedical metadata, making it a versatile tool for sample and study annotation.

## Introduction

Currently, there are millions of public omics samples available via resources like the Gene Expression Omnibus (GEO) [1], the Sequence Read Archive (SRA) [2], PRIDE [3], and MetaboLights [4]. In GEO alone, >7.1 million genomics samples from >224 thousand studies contribute to a vast collection of data from various biological contexts. This massive data collection can be incredibly valuable in revealing novel insights into the molecular basis of numerous tissues, phenotypes, diseases, and environments. However, although these data are available, finding datasets and samples relevant to a biological context of interest is still difficult because these data are described using unstructured, unstandardized, plain-text metadata, which is not easily machine-readable and unambiguously searchable [5].

To tackle this issue, significant efforts have been made to manually annotate datasets [6]. However, manual annotation is not feasible for the exponentially-growing volume of datasets, which now runs in the millions. To automate the annotation process using the metadata, natural language processing (NLP) have been employed to overcome these challenges. Rule-based NLP methods annotate metadata using text-matching or regular expressions [7, 8, 9]; however, these methods are vulnerable to misspellings or variations of a query term in the study or sample descriptions, and cannot infer annotations based on biomedical concepts in the text that are different from but relevant to a query term.

The emergence of transformer architecture-based models has revolutionized the application of NLP in the biomedical domain [10, 11]. Utilizing the power of transformer models, previous methods have framed the annotation task as translation [12] or Q&A [13]. For example, GeMI [12] uses a fine-tuned GPT-2 model to annotate a wide array of term types related to a sample, including species, sequence type, tissue, and cell type, where the “question” is metadata, and the “answer” is term types and corresponding predicted terms. Further, to combat the black-box nature of transformer models, GeMI used the saliency map technique to highlight prediction-related text. However, GPT-based models require a restructuring of the input such that it follows a fixed template, which guides the model in generating coherent and meaningful responses. This constraint makes it difficult to adapt fine-tuned models for annotating biomedical text from sources that do not fit the template. Moreover, GeMI’s output labels are not assigned to controlled vocabularies from ontologies, leading to lingering ambiguity in the annotation terms, which in turn hinders data discovery. Additionally, the GPT model is not lightweight enough to efficiently predict samples at a large scale for millions of metadata [14, 15].

Traditional machine learning (ML)-based approaches have also been used to extract information from meta-data [16, 6]. For example, Wang *et al*. [6] manually curated studies related to disease, drug, and gene perturbations from GEO. Then, they converted metadata to a term frequency-inverse document frequency (TF-IDF) matrix that considers the relative frequency of individual n-grams across all metadata documents. Combined with curated labels, these data were used to train ML classifiers to automatically extract signatures from GEO. However, since the features of the TF-IDF matrix are text, trained models cannot handle input text that includes unseen words, such as metadata from other databases, which limits the generalizability of the model.

Furthermore, the advancement of biomedical Large Language Models (LLMs) has allowed for the conversion of biomedical text to numerical representations, also called embeddings, which can effectively capture information in the text [17]. Leveraging this technique, *txt2onto* [18] represented sample descriptions as the average embeddings of words in metadata, then trained ML models to annotate tissue and cell types using these embeddings. Theoretically, the *txt2onto* framework can be adopted to annotate any kind of general biomedical text. However, one issue with the *txt2onto* approach is that averaging the embeddings of all words in a description can dampen the signal from informative biomedical terms. Furthermore, the trained model coefficients of the embedding features do not provide insight into which specific words in the sample descriptions contributed to the model’s predictions, limiting interpretability.

To tackle the challenges mentioned above, we present *txt2onto 2*.*0*, a novel and lightweight approach that assigns standardized tissue, cell type, and disease annotations to unstructured sample- and study-level meta-data. *Txt2onto 2*.*0* combines the power of semantic relationships captured by LLMs (as in *txt2onto 1*.*0* [18]) with the high interpretability offered by word-based modeling, leading to significant improvements in performance while providing transparent, easily understandable predictions. In the following sections, we demonstrate that our ML framework not only outperforms *txt2onto 1*.*0* in sample-level tissue and study-level disease annotation, especially for highly specific tissues and disease terms with limited training instances, but also highlights relevant words in each metadata associated with the annotated tissues and diseases. Moreover, our model can differentiate between similar tissues and similar diseases, enhancing the specificity of the annotations. Furthermore, we demonstrate that our disease classification models, trained on descriptions of transcirptomics studies from GEO, are proficient at infer disease annotations for biomedical metadata from any source.

## Methods

### Overview of *txt2onto 2*.*0* for disease and tissue annotation

The primary goal of our work was to develop interpretable classifiers that annotate unstructured metadata of omics samples or studies to controlled tissue, cell type, and disease terms from ontologies. To increase the interpretability of our models compared to the previous state-of-the-art method, *txt2onto 1*.*0* [18], *txt2onto 2*.*0* introduces a key improvement: instead of using average word embeddings as features, it converts sample or study metadata into a TF-IDF vector, which serves as input to the ML classifier. During the prediction phase, this classifier accepts the TF-IDF vector of each new unlabelled metadata as input and leverages word embeddings from an LLM to map words in the new metadata, including those unseen during training, to the training feature space. With text as features, our approach uses model coefficients to easily track the biomedical words/phrases that strongly influencing model predictions (**Fig. 1**). A detailed comparison of the key differences between *txt2onto 1*.*0* and *2*.*0* is provided in **Table S1**.

**Figure 1.**
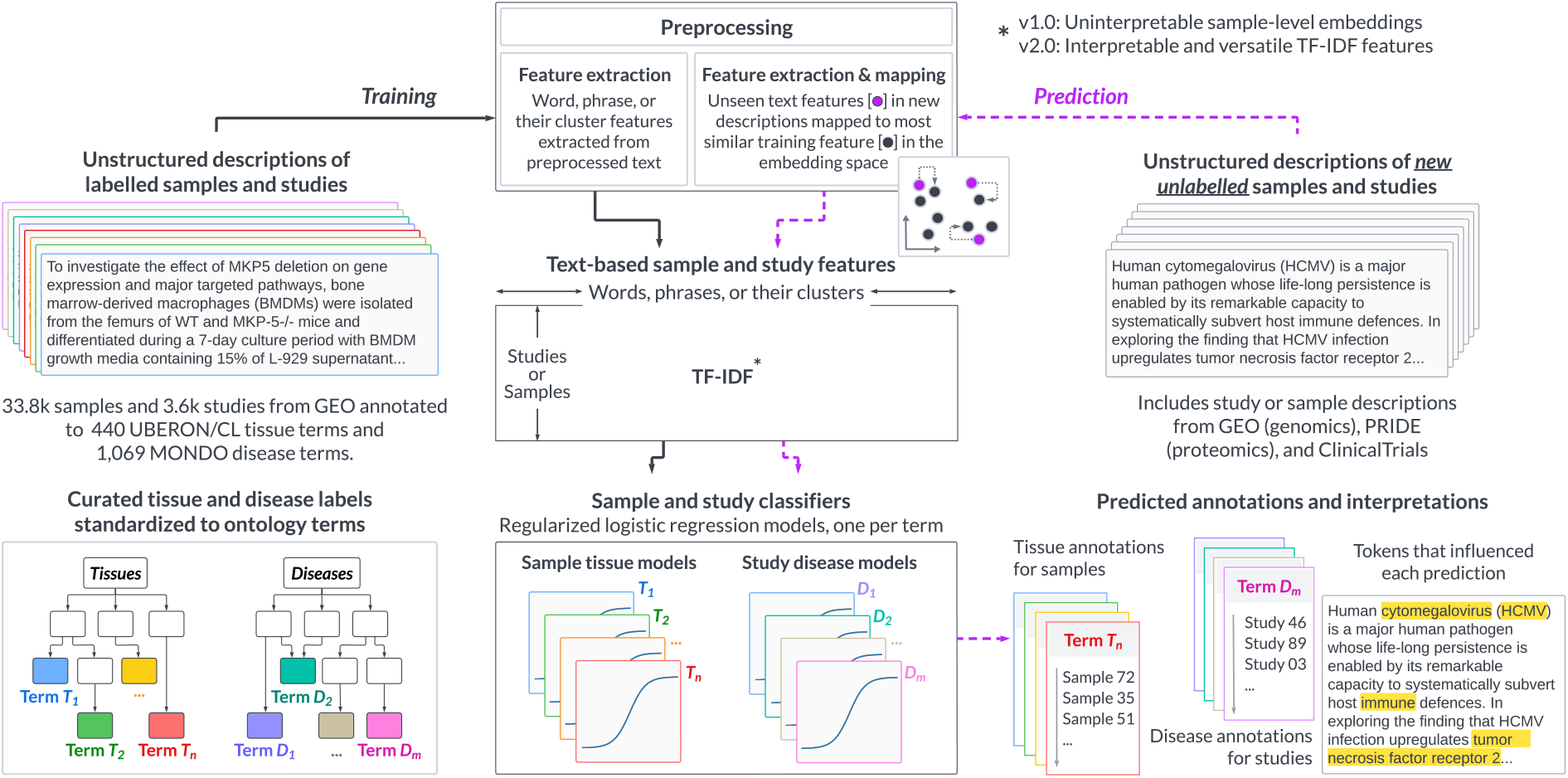
Overview of *txt2onto 2*.*0*. Samples and studies from the GEO public database are annotated with cell type, tissue, and disease ontology terms to create a gold standard dataset with 33,849 samples and 3,676 studies. The text descriptions for these samples and studies are preprocessed to remove elements that do not contribute to identifying relevant tissue or disease terms. This step generates distinct corpora for samples and studies. TF-IDF features are then derived from these corpora for various entities, including individual words, word clusters, and phrase clusters in each sample or study. A key difference between *txt2onto 2*.*0* and its predecessor is how the models generate features. Where version 1.0 used uninterpretable LLM-generated sample-level embeddings as features, version 2.0 uses interpretable TF-IDF feature vectors. A supervised classification model is trained for each tissue (n=202) or disease (n=177) term using the extracted TF-IDF features. When predicting annotations for unseen samples or studies, unlabeled text descriptions undergo the same workflow: preprocessing, TF-IDF feature extraction, and then input into the trained models, which produce a probability of a given sample belonging to a specific tissue, cell type, or disease. However, during TF-IDF feature extraction for new metadata, words or phrases that were unseen during training are mapped to their most similar word or phrase in the training feature set calculated via the cosine similarity of their embeddings.

### Collecting and processing training data

We began by collecting unstructured metadata from GEO to train the tissue and disease annotation classifiers. GEO metadata can be divided into two groups: study-level descriptions, which describe the study’s aim and design, and sample-level descriptions, which detail the source and processing of individual samples within studies. These two metadata groups were qualitatively different in terms of the tissue and disease information they contained. Sample descriptions invariably contain information about the sample’s tissue-of-origin while study descriptions often lack explicit tissue source information; further, a single study may include samples from multiple tissues. Hence, we decided to train the tissue classifiers at the sample-level. On the other hand, study descriptions contained notes about the disease being studied while sample descriptions mostly only containing modifiers such as “yes/no” or “wt/ctrl” without any mention of the disease. Therefore, we decided to train disease classifiers at the study-level.

To create the input text for these classifiers, we identified and extracted relevant fields from the original metadata. Some choices were clear: fields like source name and description contain information directly related to classification tasks while fields such as dates and contact information are irrelevant. Other fields like lab protocols and data processing methods were less clear because they may contain potentially useful information. We observed that the choice of which fields to include during training indeed impacts classifier performance (**Fig. S1**). For details about the fields used in the following analysis, see **Supplementary Methods, Preprocessing input**. After extracting the relevant fields from each metadata entry, we removed potentially uninformative elements (e.g., punctuation, URLs), converted the remaining words to lowercase, and concatenated the text from the selected fields. As we observed that lemmatization (*i*.*e*., reducing different word forms to a common root) had little to no effect on performance (**Fig. S2**), we kept all words in their original forms.

We then used the processed metadata to create a training corpus with a TF-IDF feature matrix. In this matrix, the rows represent samples or studies and the columns represent the text-based features. We extracted words and phrases (multi-word terms) as features. Further, to account for redundancy among semantically-related words and phrases, we grouped similar words or phrases into clusters, and treated each cluster as a new feature (see **Supplementary Methods, Converting preprocessed text into TF-IDF matrix**). Overall, our approach considers four types of entities as features: words, phrases, word clusters, and phrase clusters.

To train classification models, we created a gold-standard dataset consisting of 33,849 samples and 3,676 studies from various sources [19, 20, 21, 22, 18, 23, 24] (including in-house curated data; **Table S2 and S3**) labeled with terms in the Uberon-CL tissue/cell ontology [25] and the MONDO disease ontology [26]. Upon filtering for redundancy and for availability of enough examples for hyperparameter tuning and independent testing (see **Supplementary Methods, Selecting terms for model evaluation**), we were left with 202 specific tissue/cell-type terms and 177 disease terms.

### Training disease and tissue classification models

For each of the selected disease and tissue terms, using the gold-standard labels and TF-IDF feature matrix, we trained a one-vs-rest logistic regression (LR) classifier to predict whether a given sample or study is associated with that term or not. We selected LR model over other popular models such as support vector machine or random forest since LR outputs well-calibrated probabilities [27]. We used ElasticNet to regularize the classifier [28] and tuned the regularization strength and the mixing hyperparameters (see results in **Supplementary file 1**).

### Predicting labels for new samples or studies

To make a prediction for a new sample or study, its metadata text was processed and transformed into a TF-IDF vector following the same procedure as above. Given that the new metadata might contain unseen but informative entities that do not exist among the features in the training set, we mapped unseen entities to their closest training features by comparing the cosine similarity between their text embeddings from a langauge model (see **Supplementary Methods, Converting preprocessed text into TF-IDF matrix**). This approach allowed us to map the TF-IDF vector of the new metadata to the same feature space as the trained classifier. Once supplied with this input, the LR classifier for each term outputs a probability of that sample/study being annotated with that standardized tissue or disease term. Since *txt2onto 2*.*0* accepts just plain text as input, the bank of pre-trained LR classifiers can be used to make disease or tissue predictions from any type of biomedical text and are not limited to GEO metadata.

## Results

### *Txt2onto 2*.*0* outperforms *txt2onto 1*.*0* especially for terms with limited training data

We evaluated the performance of the new LR classifiers with various features (words, phrases, word clusters, and phrase clusters), comparing them to each other and to *txt2onto 1*.*0* to identify the best-performing approach (**Fig. 2**). Overall, the new models outperform *txt2onto 1*.*0* in both disease and tissue prediction tasks (**Fig. 2a**). Among the new models, those using word-based features consistently provide the best performance compared to using any of the other text features. For tissue classification, word-based and phrase-based models deliver better results compared to *txt2onto 1*.*0*, while cluster-based models fall short. For the disease classification, all our proposed feature combinations demonstrate significant improvement over *txt2onto 1*.*0*.

**Figure 2.**
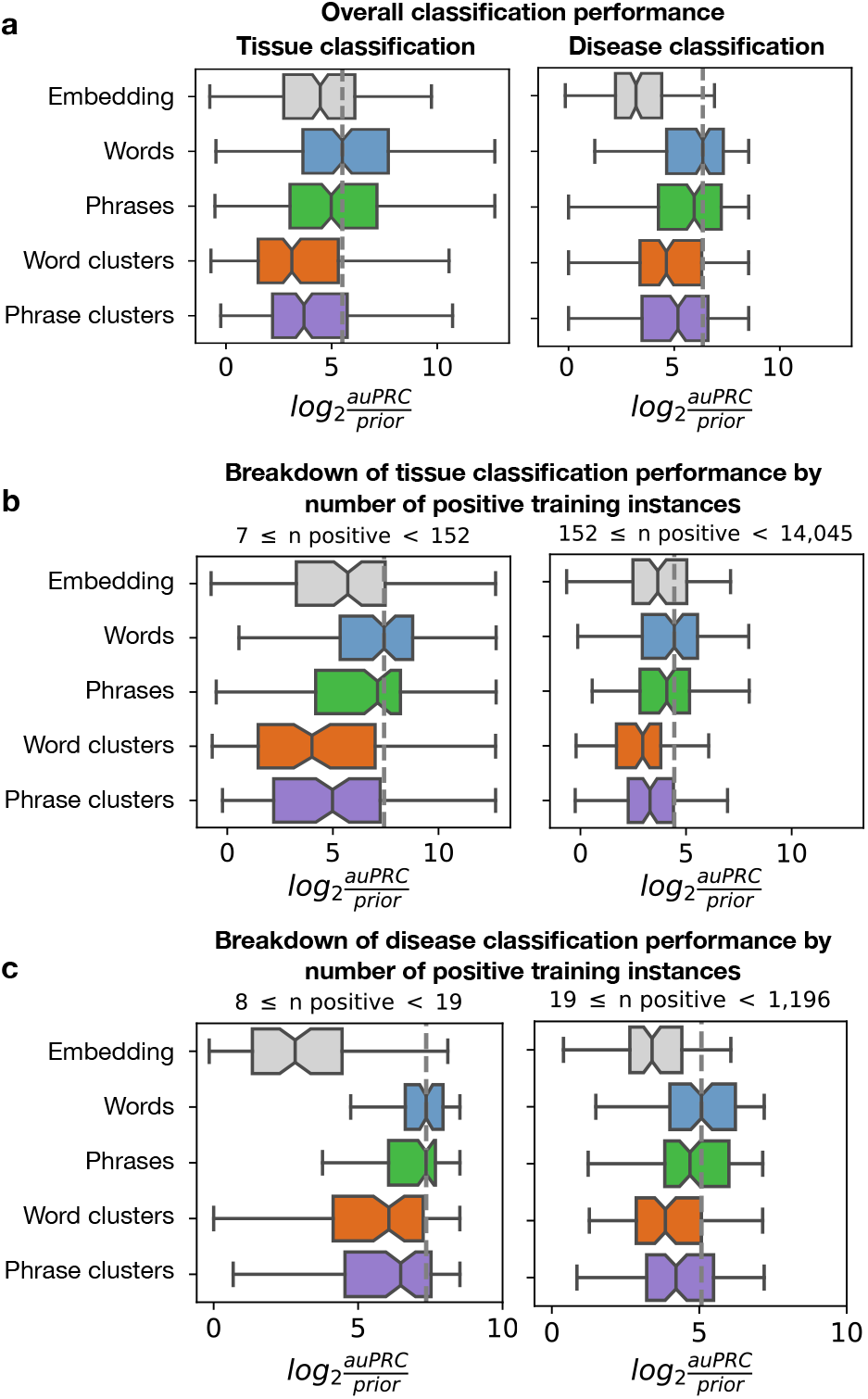
Performance of disease and tissue predictions. Boxplots display the distribution of *log*_2_(*auPRC/prior*) scores across 202 tissues and 177 diseases for each entities (words, phrases, word clusters, phrase clusters) using the LR model, along with *txt2onto 1*.*0* (embeddings). The auPRC is the area under the precision-recall curve. To account for varying class imbalances across terms, it is normalized by the prior, which is the ratio of positive instances to the total instances in the training set. Each point in the boxplots represents the performance of a disease or tissue model on the test set. Only non-redundant tasks, where the training set could be split into at least two folds, are included. The gray dashed lines indicate the median performance of the best-performing model (logistic regression with word features). (**a**) Boxplots showing overall performance for tissue (left) and disease (right) classification. (**b, c**) Boxplots in (b) and (c) show detailed performance for tissue and disease classifications, respectively, with tasks divided into two quantiles based on the number of positive instances in the training set.

Given that the number of positive instances can significantly impact model performance (**Fig. S3**), we examined the performance within subsets of terms split by the numbers of positive instances. The results show that our new models improve performance across terms with different training set sizes and are particularly suited for predicting terms when only a small number of positive examples are seen during training (**Fig. 2b-c**). This subset corresponds to terms that are either understudied in general or underrepresented in manually-curated labeled datasets. For instance, in our gold standard label set, 49 disease ontology terms contain only 8 to 11 positive instances each. We confirmed that our new model has the best performance for these terms with very few positives too (**Fig. S4a and b**). In extreme cases, some terms might have even fewer positive instances in the training set (as low as two instances), which is insufficient for hyperparameter tuning. Our model has top performance even in these cases (**Fig. S5a and b**).

Among models using different feature types, word-based models slightly outperform phrase-based models (**Fig. 2**). One reason could be the high sparsity of phrase-based feature matrices compared to word-based matrices because multi-word phrases occur less frequently than individual words. Models based on clusters of semantically-similar word and phrase features show a decrease in performance for both tissue and disease classification (**Fig. 2**). Consequently, we selected the LR model with word features as the optimal approach for all subsequent analyses.

We also investigated the potential improvement of this approach by combining predictions with MetaSRA (**Supplementary Methods, Combining *txt2onto 2*.*0* and MetaSRA predictions**). Previously, we showed that combining *txt2onto* and MetaSRA predictions leads to performance gains [18]. However, when using our novel word feature-based approach, there’s minimal performance improvement when combining tissue predictions from both models (**Fig. S6**).

This choice was also confirmed by the comparison of the LR model with an even simpler baseline model that makes predictions by greedily aggregating the occurrence individual features one-by-one, weighted by each feature’s informativeness (**Supplementary Note 1**). Although the performance of this baseline method is comparable to LR, it is oversensitive to uninformative words (**Fig. S7 and S8**) and cannot output probabilities.

### *Txt2onto 2*.*0* models learn features relevant to tissue and disease classification tasks

One of the primary goals in this new version of *txt2onto* was to develop interpretable models that capture words and phrases in sample and study metadata that are most predictive of annotations to specific ontology terms. To inspect the interpretability of the models, we summarized predictive word features from topperforming tissue and disease models as word clouds, where the size of each is proportional to its regression coefficient. For example, the keywords for *Glucagon-secreting cells* (CL:000170) (**Fig. 3a top**) are “alpha”, “islets”, “pancreatic”, “developmental”, “fetal”, “adult”, “stage”, and “gestational”. These terms refer to the endocrine function of the pancreas, with “alpha” referring to alpha cells that produce glucagon, a hormone that regulates blood sugar levels. “Islets” are the islets of Langerhans, clusters of cells in the pancreas. The words “developmental” and “gestational” imply a context of fetal (early developmental) or adult stages where these cells are involved. The informative words for *Coronary artery disorder* (MONDO:0005010) (**Fig. 3b top**) include “coronary”, “artery”, “disease”, “stenosis”, “atherosclerosis”, “myocardial”, “CAD”, “heart”, “blood”, and “ventricular”. These words are clearly linked to the cardiovascular system, specifically the arteries that supply blood to the heart. Coronary artery disease (CAD) involves the narrowing of the arteries due to atherosclerosis, potentially leading to myocardial infarction or heart attacks. Similarly, the appearance of key terms such as “MECP2” for *Rett syndrome* (**Fig. 3b middle**), “merlin” for *Meningioma* (**Fig. 3b bottom**), “sickle” for *Erythrocyte* **(Fig. 3a middle**), and “bronchoalveolar” for *Pulmonary acinus* (**Fig. 3a bottom**) demonstrate that our models successfully identify relevant features from the metadata text that significantly contribute to its the correct tissue and disease annotation predictions.

**Figure 3.**
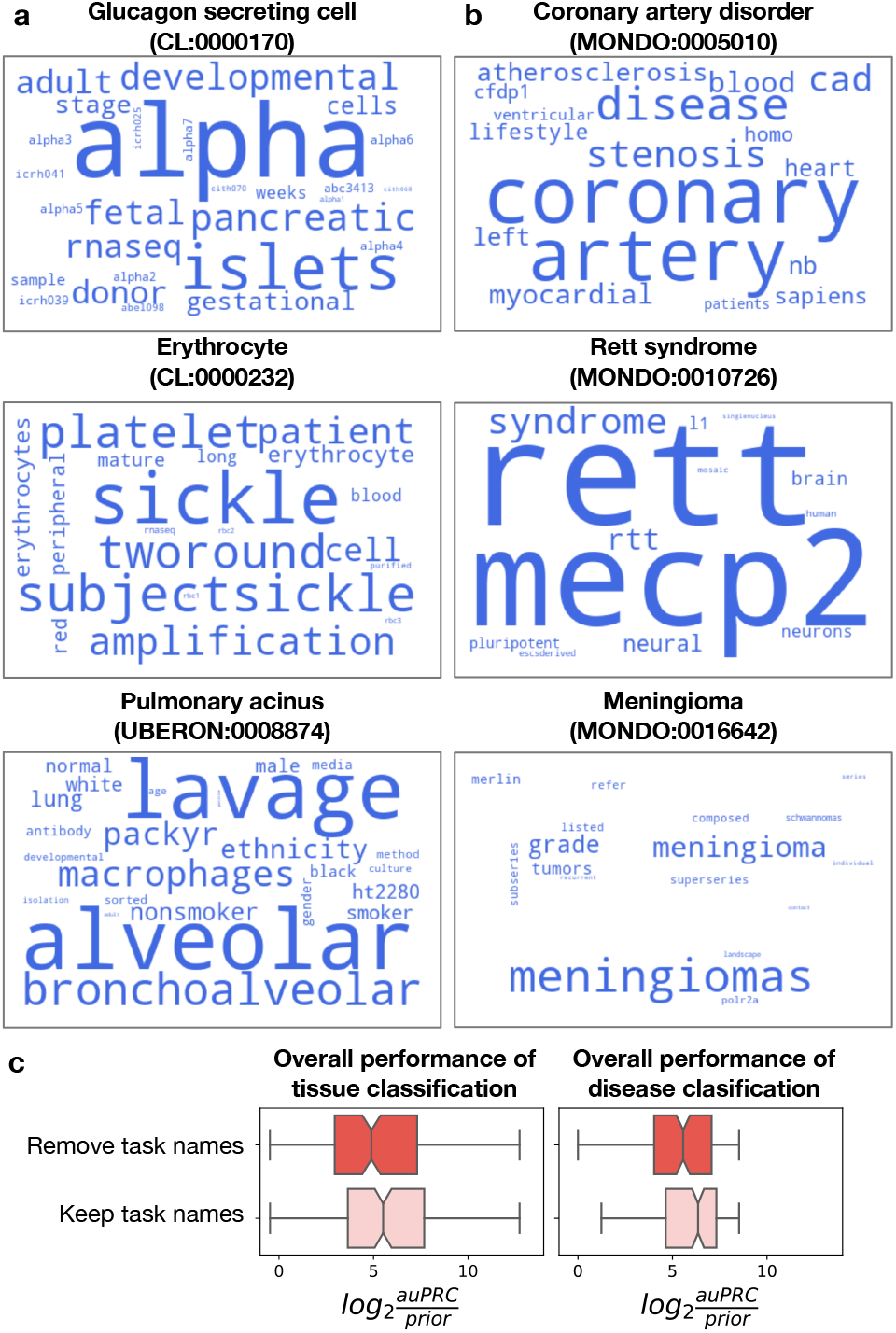
Predictive words for disease and tissue classification. (**a, b**) Word clouds highlight the most predictive word features in the top-performing three tissue (a) and disease (b) classification models, which were trained using the best-performing model (logistic regression with word features). The size of each word is proportional to its regression coefficient in the logistic regression model, with larger words indicating higher coefficients and greater predictive importance. (**c**) The boxplots compare the change in the distribution of *log*_2_(*auPRC/prior*) scores for tissue (left) and disease (right) classification performance when task names are removed from the corpus.

Worthy of note is that, by leveraging their ability to capture term-related words and phrases, our models are able to annotate samples/studies even if the name of the tissue or disease term is absent in the metadata. *Pulmonary acinus* (UBERON:0008874) (**Fig. 3a**) is one among many good examples of terms to which where metadata are correctly annotated solely via the words and phrases associated with that term. To demonstrate this potential across models for both tissue and disease classification tasks, for each term, we compared the performance of our models before and after removing every mention of that term’s name from the metadata corpus. Results show that removing the ontology term names leads to a negligible drop in performance, demonstrating our models’ robustness to missing information in metadata (**Fig. 3c**).

### *Txt2onto 2*.*0* models accurately predict instances related to specific tissues and diseases

To truly promote the discovery of existing data relevant to specific biomedical questions of interest, it is crucial that metadata annotation is accurate for specific tissue and disease terms (e.g., *Rett syndrome*) and not just general terms (e.g., *Neurological disorder*). Therefore, we evaluated the effect of term specificity on *txt2onto 2*.*0* ‘s disease and tissue annotation performance. Using the structure of the appropriate ontology the tissue and disease term are a part of, we defined each term’s specificity using its information content (IC) of a term. Higher IC indicates specific terms closer to the leaf nodes of the ontology while lower IC indicates general terms closer to the root of the ontology. We included all disease and tissue terms without redundancy filtering in the analysis to help examine model performance across a wide range of term specificities.

This analysis showed that there is a clear association between tissue/disease term specificity (IC) and model performance (**Fig. 4**). The smaller data points, representing terms with fewer positive samples in the training set, tend to cluster towards the higher end of the information content scale and the performance metric. When cast in the context of the number of positive examples available during training, these trends indicate that our models are capable of accurately annotating to very specific terms in both the tissue and disease ontologies even when the amount of training data is limited. This trend persists across models trained on various features, including phrases, word embeddings, phrase clusters, and word clusters (**Fig. S9**). When we examined four tissue and four disease outlier models which are outliers of this trend, i.e., specific (high IC) terms having low performance (**Supplementary Note 2**), we found that poor performance often resulted from mislabeled samples or studies, suggesting that tasks with very few positive instances in the training data are sensitive to mislabeling.

**Figure 4.**
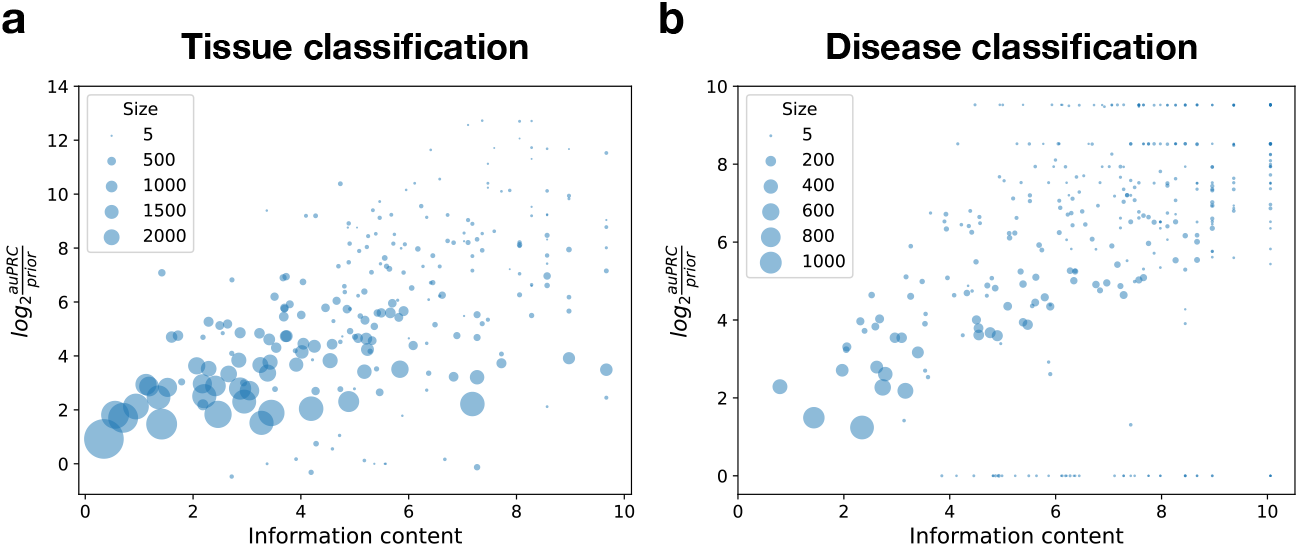
Effect of term specificity on prediction performance. Scatter plots show the prediction performance of the best model (logistic regression with word features) for tissue (**a**) and disease (**b**) classification tasks across various levels of term specificity. Each point represents the performance of a single tissue or disease model. The x-axis shows the specificity of the disease or tissue terms, defined by their information content (IC). The IC of a term is the negative logarithm of the fraction of terms in the ontology that are descendants of term of interest. The y-axis shows the model performance, measured by *log*_2_(*auPRC/prior*). The size of each data point is proportional to the number of positive instances in the training set for each task.

### *Txt2onto 2*.*0* models can differentiate between similar tissues or diseases

Following the specificity analysis, we evaluated the ability of the *txt2onto 2*.*0* models to correctly differentiate between similar or related diseases or tissues, such as distinguishing *Crohn’s disease* from *Ulcerative colitis*, or *Skeletal muscle* from *Smooth muscle*. For this evaluation, we defined the semantic similarity between pairs of terms within an ontology as the cosine similarity between their corresponding text embedding vectors calculated using BioMedBERT [29] (see **Supplementary Methods, Differentiate similar terms**). Then, we split all term pairs based on the cosine similarity into four equal-size quantile bins corresponding to term pairs with increasing similarity. Finally, within each bin, we quantified (using area under the receiver operating characteristic curve; auROC) each term model’s ability to assign higher probabilities to instances (sample or studies) labeled to that term compared to instances labeled to other terms in the same bin.

The auROC distribution within each bin being narrow and centered close to 1.0 confirms that our models can differentiate between terms irrespective of their semantic similarity (**Fig. 5**). Particularly, the *txt2onto 2*.*0* classifiers can, in general, correctly prioritize instances annotated to the correct term over instances annotated to a closely-related term. However, as reflected in the increased number of outliers in the auROC distributions, there are indeed cases where highly semantically-similar terms prove challenging for our models. We examined four such challenging disease pairs, aiming to understand why they are difficult to distinguish (**Supplementary Note 3**). The prominent reason seems to be that predictive words extracted from the input text overlap between the disease pairs (**Fig. S10**). For instance, the words “lung”, “cell”, and “cancer” appeared frequently in the metadata of instances annotated to *Squamous cell lung carcinoma* and *Small cell lung carcinoma* (**Fig. S10e**). Although “squamous” is a predictive word for *Squamous cell lung carcinoma*, this word does not appear frequently among studies of this disease. Consequently, the signal from the word “squamous” is not strong enough to be useful in effectively distinguishing between these two textually-similar lung cancers. We observed that such dilution or absence of discerning predictive words in metadata might result from mislabeling. For example, GSE28582 and GSE21933, two studies annotated to *Squamous cell lung carcinoma* in our current gold-standard, should instead have been appropriately labeled as *Non-small cell lung carcinoma*, a more general disease term, which would explain the absence of the word “squamous” in the metadata of these two studies.

**Figure 5.**
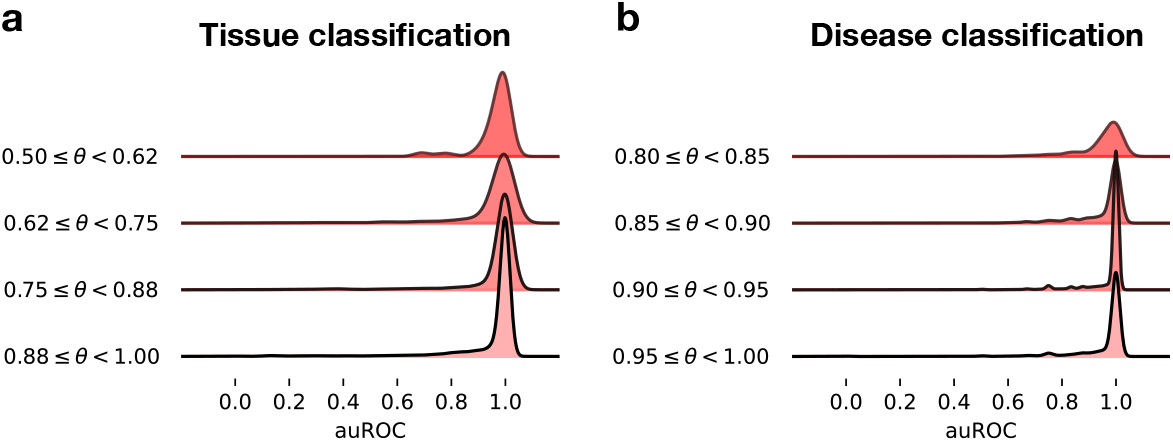
Model’s ability to annotate to the correct disease or tissue term over similar terms. Ridgeline plots show the distribution of auROC scores for the best-performing model (logistic regression with word features) in distinguishing samples from pairs of (**a**) tissue terms and (**b**) disease terms with varying levels of semantic similarity (*θ*). Semantic similarities between terms are measured by the cosine similarity of text embeddings, where larger values indicate greater semantic similarity. The range of semantic similarities, from minimum to one, is divided into four equal-sized intervals. The x-axis represents the auROC score, while the height of each ridgeline plot represents the frequency of term pairs at each auROC level.

### *Txt2onto 2*.*0* models accurately predict disease labels for completely independent studies

Although all the models were only trained on metadata from GEO (principally from transcriptomics studies), we wanted to test the generalizability of the models by evaluating their ability to annotate the metadata of studies from completely independent data repositories. As study-level descriptions are more widely available than tissue-level descriptions, we focused on evaluating the disease annotation performance. First, we assessed if our models trained on transcriptomics metadata could annotate metadata of other omics studies. For this analysis, we used our models to annotate proteomics studies from the Proteomics Identification Database (PRIDE) [3]. Second, we tested whether our models could annotate biomedical studies beyond omics. We used the descriptions of clinical trials across various biomedical domains from ClinicalTrials [30] as a case study for this test. To quantify the prediction performance on these independent datasets, we conducted a double-blind test to manually assess the predictions from the top six models (see **Supplementary Methods, Predicting on independent dataset**). Our models achieved high accuracy and F1 scores when predicting disease labels for studies from PRIDE (**Table 1**) and ClinicalTrials (**Table 2**), demonstrating the potential generalizability of *txt2onto 2*.*0*.

**Table 1.** Disease annotation performance for studies from PRIDE. This table displays the annotation performance of the top six disease classification models, evaluated based on their top ten and bottom ten predictions. Each row corresponds to a disease term, and each column represents a performance metric. The metrics included are: Acc (Accuracy), calculated from unshuffled predictions; log2(Acc/R Acc), the accuracy normalized by the random accuracy calculated from shuffled predictions, then log base two transformed; and log2(F1/R F1), the F1 score normalized by the random F1 score calculated from shuffled predictions, then log base two transformed.

| Term | Acc | F1 | log2(Acc/R Acc) | log2(F1/R F1) |
| --- | --- | --- | --- | --- |
| Invasive ductal breast carcinoma (MONDO:0004953) | 0.83 | 0.67 | 1.30 | -0.76 |
| Cytomegalovirus infection (MONDO:0005132) | 1.00 | 1.00 | 1.18 | 1.24 |
| Pancreatic ductal adenocarcinoma (MONDO:0005184) | 0.87 | 0.83 | 0.15 | 0.22 |
| Uveal melanoma (MONDO:0006486) | 0.77 | 0.68 | 0.89 | 4.26 |
| Head and neck squamous cell carcinoma (MONDO:0010150) | 1.00 | 1.00 | 1.29 | 1.29 |
| Covid-19 (MONDO:0100096) | 1.00 | 1.00 | 1.29 | 1.29 |

**Table 2.**
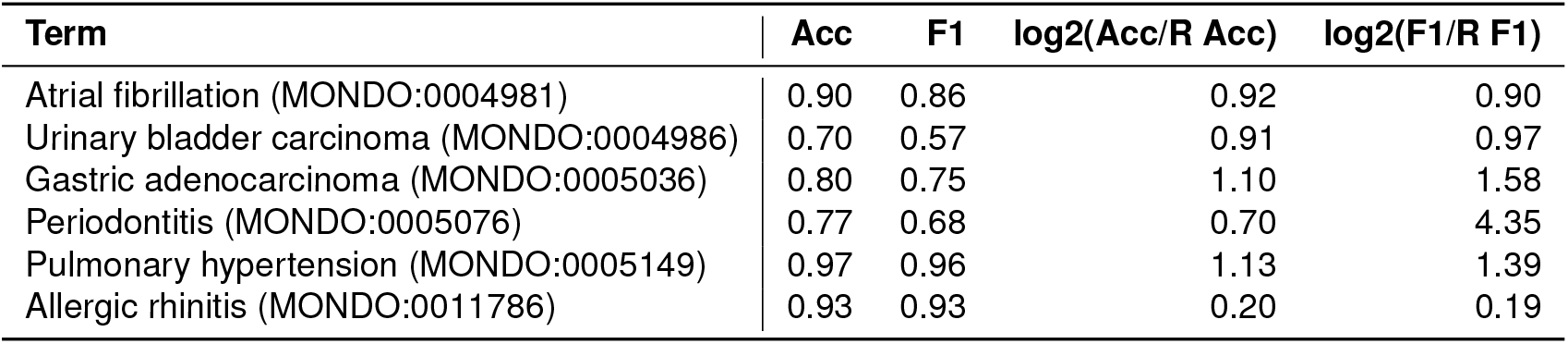
Disease annotation performance for studies from ClinicalTrials. This table presents the annotation performance of the top six disease classification models, evaluated based on their top ten and bottom ten predictions. Each row corresponds to a disease term, and each column represents a performance metric. The metrics included are: Acc (Accuracy), calculated from unshuffled predictions; log2(Acc/R Acc), the accuracy normalized by the random accuracy calculated from shuffled predictions, then log base two transformed; and log2(F1/R F1), the F1 score normalized by the random F1 score calculated from shuffled predictions, then log base two transformed.

Inspecting the predictive words in the input metadata confirmed that the text signals in these independent datasets that drive specific disease annotations are indeed related to the disease (**Fig. 6**). For example, the annotation of ClinicalTrials studies to *Allergic rhinitis* (MONDO:00117886) is based on words like “allergic”, “rhinitis”, “nasal”, “allergen”, and “asthma”. These words align with typical symptoms and triggers for allergic rhinitis, which involves nasal inflammation due to allergens (**Fig. 6a**). Similarly, the annotation of PRIDE (proteomics) studies to *COVID-19* (MONDO:0100096) is based on keywords such as “sarscov2”, “covid19”, “infection”, “severe”, and “sarscov”. These words reflect the main factors associated with COVID-19, including its causative virus, SARS-CoV-2, which leads to infection (**Fig. 6b**). Another noteworthy example is the prediction to the term *Gastric adenocarcinoma* (MONDO:0005036). Here, the word “gastrectomy” is present in metadata of relevant ClinicalTrials (**Fig. 6a**) studies influences their annotation, but is not one of training features for this disease. Nevertheless, our model aligns this unseen word to its closest training feature “gastric”, enabling it to contribute to the prediction. Examples like these affirm our approach’s capability of effectively using even previously unseen words if they are related to the annotation task under consideration.

**Figure 6.**
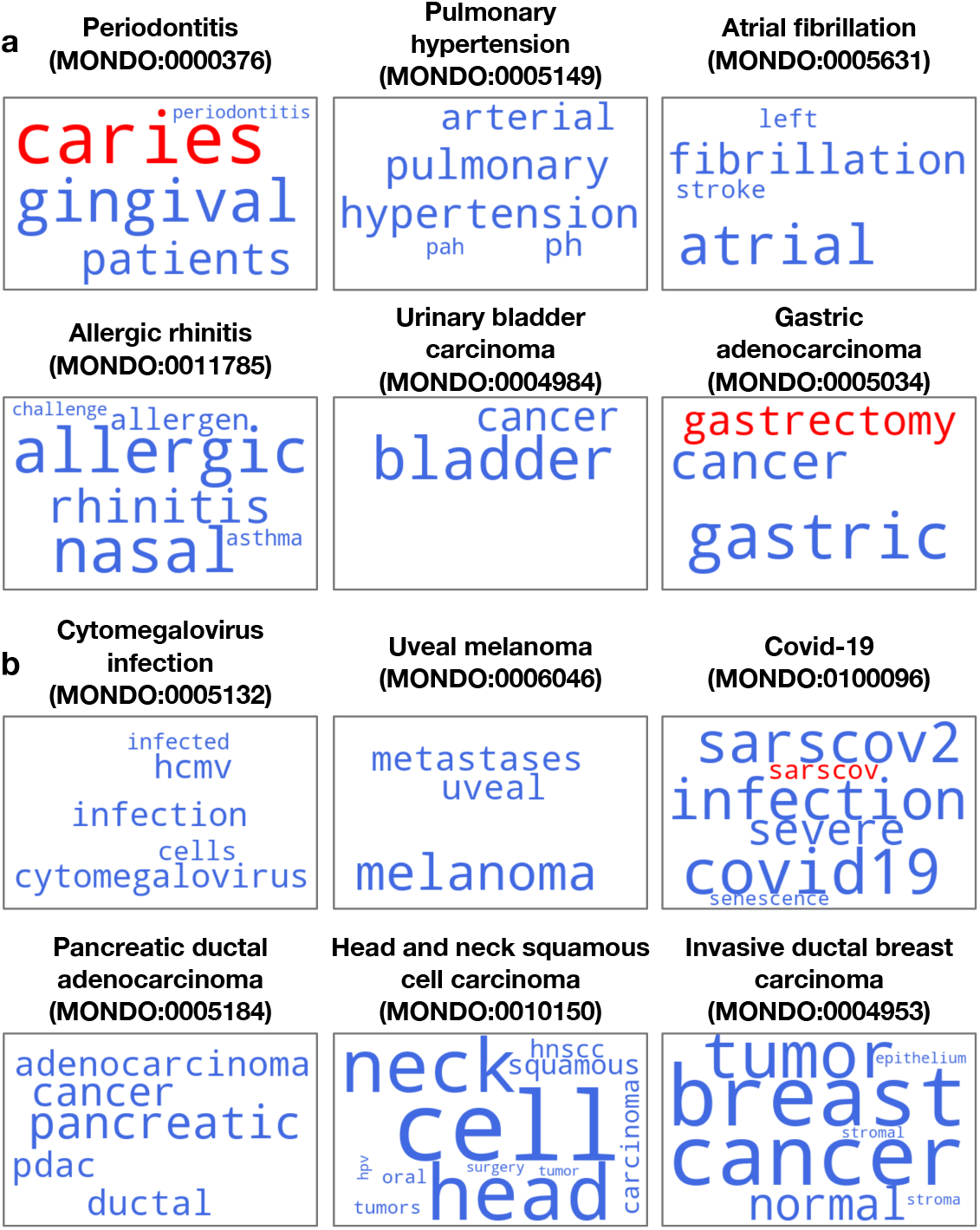
Disease-related words derived from databases containing information beyond expression profiles. Word clouds display disease-related words extracted from study titles and summaries of two independent datasets: (**a**) ClinicalTrials and (**b**) the PRIDE database. These predictions were generated by disease models trained on metadata from the GEO database, using the best-performing model (logistic regression with word features). The size of each word is proportional to its frequency among disease-related words in ClinicalTrials or PRIDE, with larger words being more prevalent. Blue words appear in both the predictive words from the independent datasets and the training features of the models, while red words are unique to the independent datasets.

More detailed results for the performance of our models in annotating text data from data sources other than GEO are given in the supplementary materials (see **Table S4-S5**). Together, these analyses demonstrate the ability of *txt2onto 2*.*0* in annotating any biomedical text description to appropriate ontology terms. The interpretability of our models allows one to trace back the words contributing to the prediction, providing a way to easily evaluate and confirm the annotations.

## Discussion and Conclusion

Reusing existing biomedical data is paramount to advancing scientific research and accelerating medical discoveries [31, 32]. However, the samples and studies stored in current vast biomedical data repositories are often described using unstandardized, unstructured, plain-text descriptions. This poor quality of metadata is a major hindrance for researchers in discovering the datasets most relevant to their context or question of interest. In this work, we propose *txt2onto 2*.*0*, a computational approach that combines NLP techniques and ML to annotate any biomedical descriptions to standardized tissue and disease ontology terms. Thus, by providing a way to index public metadata for easy search using controlled vocabularies, *txt2onto 2*.*0* addresses the first goal of the FAIR (Findable, Accessible, Interoperable, and Reusable) principles [33].

Through systematic and rigorous evaluations, we have demonstrated that our method outperforms the state-of-the-art method, *txt2onto 1*.*0* [18], in disease and tissue classification. Especially in disease classification, our method markedly outperforms *txt2onto 1*.*0*, particularly when dealing with highly imbalanced cases. Disease classification poses greater challenges than tissue classification for the previous version due to the longer study descriptions used to infer diseases (**Fig. S11**). Long text inputs are not amenable to *txt2onto 1*.*0* ‘s idea of representing metadata as the average of the embeddings of all the constituent words, which dampens the signal from informative words. *Txt2onto 2*.*0* overcomes this limitation by utilizing a TF-IDF feature matrix to represent text, naturally avoiding the mixing of signals from predictive words with the rest of the text. This strategy enables *txt2onto 2*.*0* to predict a wider range of tasks than *txt2onto 1*.*0*, allowing for accurate predictions of understudied tissues and diseases.

In addition to inferring accurate tissue and disease annotations of metadata, it is crucial to address interpretability and explainability to increase trust in and verifiability of the annotation results. By using word-level features combined with a large language model, *txt2onto 2*.*0* achieves both these desired qualities: i) Each tissue or disease term model is highly interpretable in terms of the most informative text features learned during training, and ii) Each predicted annotation (of an input metadata to a particular tissue or disease term) is explainable in terms of the specific text snippets in the new metadata that drove the prediction. As a contrast, using average word embeddings as features makes in *txt2onto 1*.*0* makes both interpretability and explainability infeasible. Recent GPT-based models are promising for extracting disease and tissue labels from unstructured text due to their strong text comprehension powers. However, they operate as “black boxes”. Efforts in explainable AI, such as GeMI’s use of saliency maps to highlight words related to an annotation, have been made to explore the reasoning behind such models [13]. Nevertheless, these techniques have limitations, such as providing only post-hoc explanations without deeper insights into the model’s internal logic and lacking systematic evaluation of interpretability. In the era of LLMs, we acknowledge that using better models like GPT-4 and state-of-the-art explainable AI methods could achieve better interpretability and explainability. However, it is critical to continue exploring simple and elegant methods like *txt2onto 2*.*0* for building models with high annotation accuracy and inherent transparency that are also lightweight and cost-effective so that they can scale to exponentially growing number of samples and studies across biomedical data repositories.

Extremely imbalanced data is a common challenge in biomedical prediction studies [34] — an issue also present in ours (**Fig. S3**) due to most annotation tasks (tissue and disease terms) only having limited number of positive instances. Despite using a LR model, which typically outputs well-calibrated probabilities, the high class imbalance leads to underestimated probabilities for positive instances. We applied two strategies to identify likely positive predictions while dealing with underestimated probabilities: (1) Train a recalibration model to project underestimated probabilities to the 0–1 range with pivot point close to 0.5, or (2) compare predicted probabilities to expected random classification performance. We did not choose undersampling or oversampling strategies as they might not work for extremely imbalanced datasets, and correcting class imbalance may worsen the reliability of estimates [35]. Additionally, we found that models trained and tested on imbalanced datasets are extremely vulnerable to mislabeled instances because incorrect signals can easily dominate training or evaluation, leading to poor performance (see **Supplementary Note 2**). Possible approaches to combat this problem include using methods that allow for naturally updating the labels, such as active learning [36] and semi-supervised learning [37], or building a unified model for all tissue or disease classification tasks to borrow information from related tasks.

Given that inferring standardized annotations of metadata is a universal problem pertinent to discovering and reusing data in every biomedical repository, generalizability is a fundamental requirement of automatic annotation inference methods. It is important to realize this generalizability without having to (re-)training the underlying models based on curated metadata labels from every database. The models also need to be versatile in dealing with any generic biomedical metadata, regardless of the varying format or writing style. Our method meets this need by treating metadata as a bag-of-words, making it flexible enough to accept input from any source. Further, by using the latent embedding space that captures semantic similarities between words, *txt2onto 2*.*0* is able to utilize words in new metadata to inform an annotation even if those words were never seen during training (**Fig. 6a**).

Our study opens up numerous avenues for refinement and exploration in the future. First, the informative keywords identified by our method occasionally include generic words because they are overrepresented among the few positive instances (compared to negative instances) available during training. For instance, words such as “sample”, “week”, and “cells” are highlighted as predictive words for *Glucagon secreting cell* (CL:0000170) (**Fig. 3a**) despite not being specifically related to this cell type. This problem could be more prevalent in tissue annotation (of samples) compared to disease annotation (of studies), as the richer context in study summaries reduces the likelihood of selecting spurious words, while the limited context in sample descriptions makes the model more prone to latching on to generic words. In future work, we may employ causal inference techniques [38] or map extracted terms to biomedical knowledge graphs and perform graphbased reasoning to remove irrelevant words [39]. Second, we annotated disease labels at the study level but did not attempt sample-level disease annotation, *i*.*e*., annotating samples within a study as ‘healthy’ and ‘disease’. This is because sample-level metadata in public databases is severely lacking in completeness and quality (*e*.*g*., sample metadata lacking any information about the disease being studied and containing, if at all available, only indicators like ‘yes’, ‘wt’, or ‘ctrl’). In the future, sample-level disease annotation could be achieved by employing additional computational models that integrate information from samples across studies to train a single model to classify between healthy and disease samples. Third, future work can expand to other annotation categories beyond tissue and disease such as sex, age, phenotype, or environment. Finally, moving from one model per term towards a unified model for annotating multiple terms via a shared representation learning will likely lead to improve prediction performance across the board, and especially for terms with few positive instances.

Overall, *txt2onto 2*.*0* is a novel, light-weight, interpretable, and explainable ML-based approach to annotate biomedical text from various sources with standardized tissue and disease labels, even when there are limited amounts of training instances. Biomedical researchers will be able to use the labels predicted by our method to drastically improve data organization and curation, and to effectively reuse existing data to make potentially novel scientific discoveries in downstream analyses. To ensure that our approach can benefit the research community for data reuse, we released *txt2onto 2*.*0* on our GitHub repository. Users can either predict disease or tissue labels using provided models or build their own models from scratch.

## Code and Data Availability

We have made the trained disease and tissue classification models, a Python script for classification, and demo scripts, along with extensive documentation available at https://github.com/krishnanlab/txt2onto2.0 (v1.0.0). The repository also includes utilities for training new models for a user-defined task.

## Acknowledgement

We would like to thank Keenan Manpearl for valuable suggestions on the organizations and documents of the Github repository, as well as code testing. We also would like to thank all members of the Krishnan Lab for valuable discussions and feedback on the project.

## Funding

This work was primarily supported by the National Science Foundation grant 2328140 to A.K.

## Author contributions

H.Y. and A.K. designed the study. H.Y. developed the software. H.Y., K.J. and L.V. curated annotations. H.Y., P.H., M.A. and K.J. performed the analyses. H.Y., P.H., M.A. and K.J. interpreted the results. H.Y., P.H. and M.A. wrote the final manuscript with feedback from A.K.

## Author Biographies

**Hao Yuan** is a Ph.D. candidate at Michigan State University. His research focuses on applying natural language processing techniques to biomedical metadata and leveraging network biology approaches for personalized medicine.

**Parker Hicks** is a Ph.D. student at the University of Colorado Anschutz Medical Campus. His work is centered around natural language processing of biomedical metadata and analysis of transcriptomics profiles.

**Mansooreh Ahmadian** is a Research Associate in the Department of Biostatistics and Informatics at the School of Public Health, University of Colorado Anschutz Medical Campus. Her research focuses on developing machine learning-based analytical pipelines to extract knowledge from biomedical and clinical data.

**Kayla Johnson** was a postdoctoral fellow at the University of Colorado Anschutz Medical Campus. Now, she is a computational biologist at Synthesize Bio. Her research primarily focuses on computational methods for human health and diseases.

**Lydia Valtadoros** is a research assistant at the University of Colorado Anschutz Medical Campus. Her research currently focuses on biological sample metadata and sex bias in biomedical research areas.

**Arjun Krishnan** is an associate professor at the University of Colorado Anschutz Medical Campus. His group works broadly in the areas of computational genomics and biomedical data science.

## Notes

### Competing Interest Statement

The authors have declared no competing interest.

### Summary of Updates

We added another analysis for combining txt2onto 2.0 results with metasra, and access performance improvement. We also added detail in figure 1 about the difference between txt2onto 2.0 and 1.0.

