## Supplementary material for "Annotating publicly-available samples and studies using interpretable modeling of unstructured metadata"

#### Contents


**Supplementary Figures** **2**

**Supplementary Tables** **14**

**Supplementary Methods** **17**

**Supplementary Note 1** **21**

**Supplementary Note 2** **22**

**Supplementary Note 3** **23**

**Supplementary Note 4** **25**

### Supplementary Figures

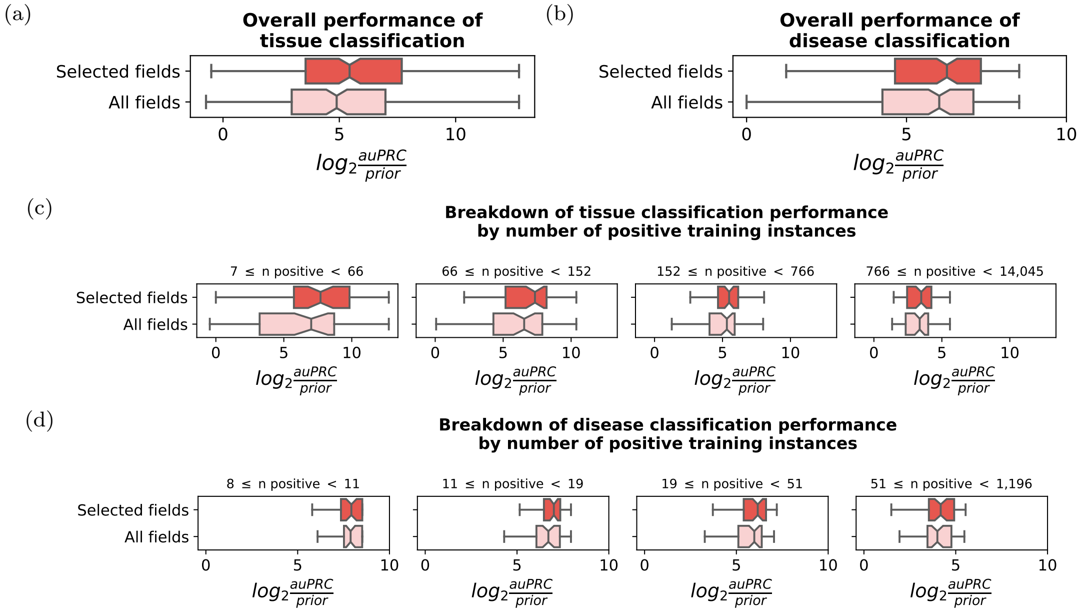


Figure S1: **Effect of task-irrelevant fields on model performance.** The boxplots show the effect of removing task-irrelevant fields on tissue and disease classification performance, measured by $\log_2$(auPRC/prior). The predictions are made using the best-performing model (logistic regression model with word features), while the input corpus might only include selected fields (top) or keep all fields (bottom). (**a, b**) The boxplots show the effect of removing task-irrelevant fields on the overall performance of tissue (a) and disease (b) classification. (**c, d**) The boxplots show the breakdown of tissue (c) and disease (d) classification performance. Tasks are divided into four quantiles with an equal number of tasks based on the number of positive instances in the training set.

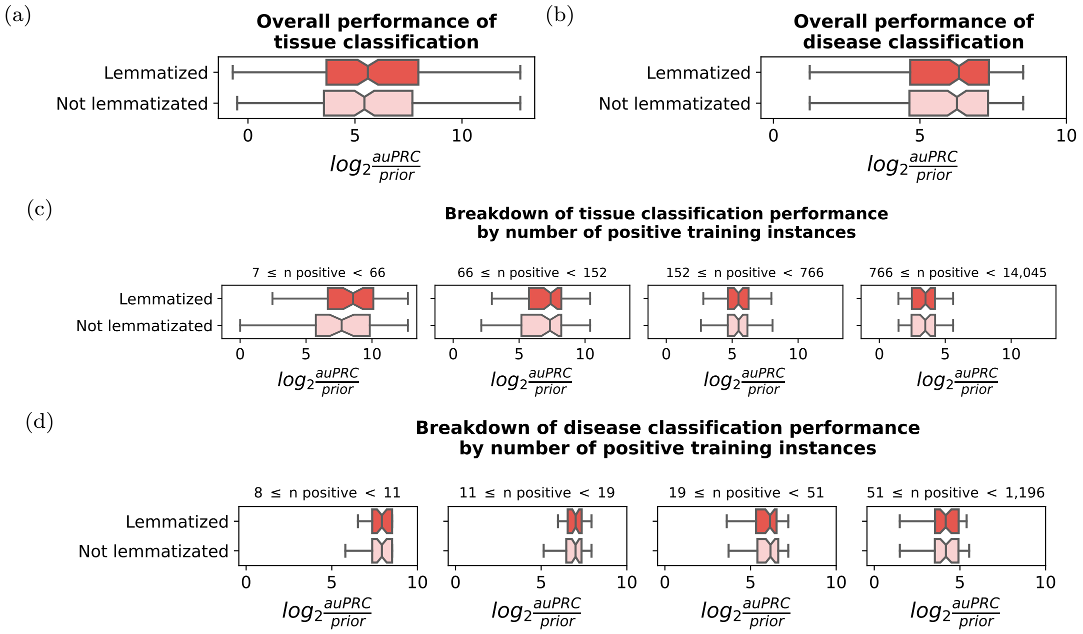


Figure S2: **Effect of lemmatization on model performance.** The boxplots show the impact of lemmatization on the predictive performance, measured by $\log_2$(auPRC/prior). The predictions are made by the best-performing model (logistic regression model with word features), while the input corpus might be lemmatized (top) or unprocessed (bottom). (**a, b**) The boxplots show the effect of lemmatization on the overall performance of tissue (a) and disease (b) classification. (**c, d**) The boxplots show the breakdown of tissue (c) and disease (d) classification performance. Tasks are divided into four quantiles with an equal number of tasks based on the number of positive instances in the training set.

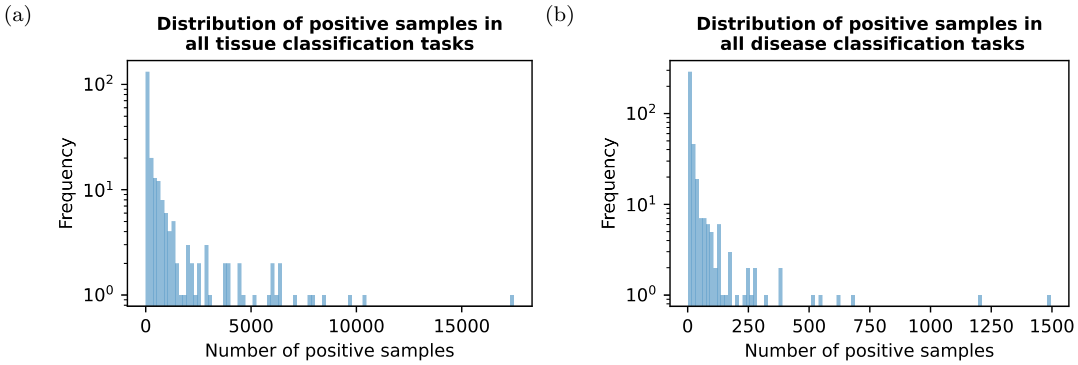


Figure S3: **Distribution of the number of positive instances among gold standards.** The histograms show the distribution of the number of positive instances in all tissue (**a**) and disease (**b**) tasks from the gold standards. The labels of instances in the gold standards have been propagated to general terms. The frequency on the y-axis is displayed on a logarithmic scale.

(a)

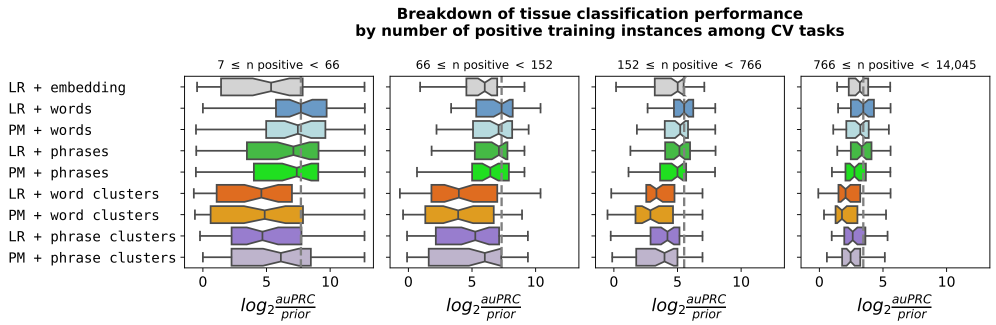


(b)

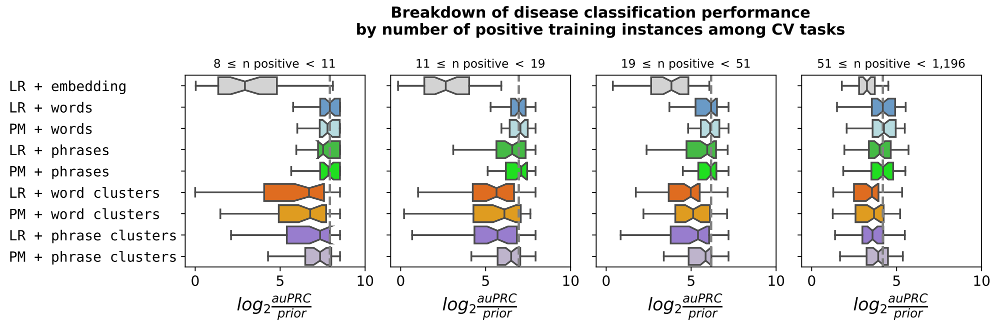


Figure S4: **Breakdown of tissue and disease classification performance on non-redundant tasks for hyperparameter tuning.** The boxplots show the $\log_2$(auPRC/prior) scores for the breakdown of tissue (**a**) and disease (**b**) classification performance for each of the entities (words, phrases, word clusters, phrase clusters) with logistic regression (LR) and power matrix (PM) model, as well as *txt2onto 1.0* (LR + Embedding). Only non-redundant tasks, where the training set could be split into at least two folds, are included. Each point in the boxplots corresponds to the performance of a single tissue or disease model. The grey dashed line represents the median performance of the best-performing model (LR + words). Tasks are divided into four quantiles with an equal number of tasks based on the number of positive instances in the training set.

(a)

(b)

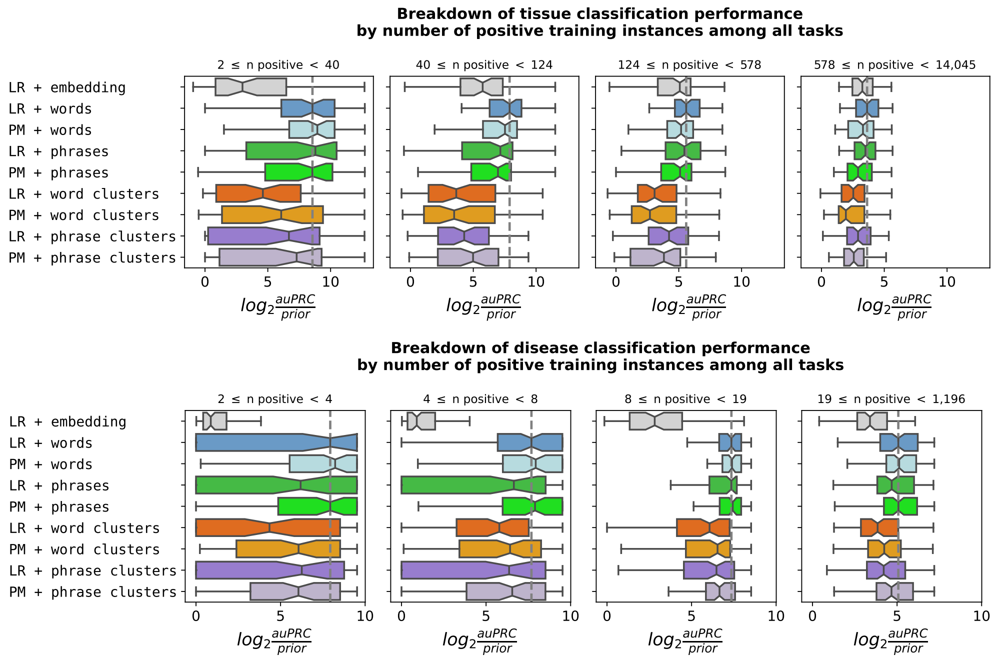


Figure S5: **Breakdown of tissue and disease classification performance on non-redundant tasks for all trainable tasks.** The boxplots show the $\log_2$(auPRC/prior) scores for the breakdown of tissue (**a**) and disease (**b**) classification performance for each of the entities (words, phrases, word clusters, phrase clusters) with logistic regression (LR) and power matrix (PM) model, as well as *txt2onto 1.0* (LR + Embedding). All non-redundant tasks are used to assess the performance, as long as they can be divided into training and testing sets. Each point in the boxplots corresponds to the performance of a single tissue or disease model. The grey dashed line represents the median performance of the best-performing model (LR + words). Tasks are divided into four quantiles with an equal number of tasks based on the number of positive instances in the training set.

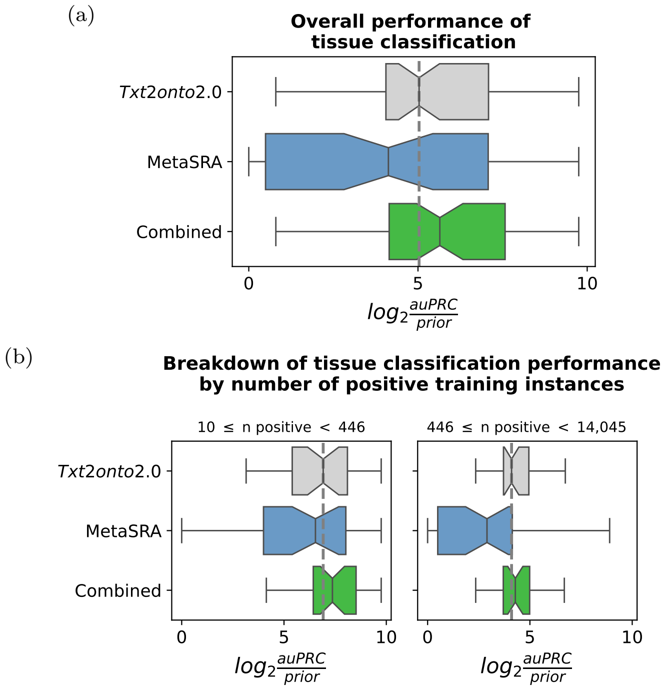


Figure S6: **Tissue classification performance when combining *txt2onto 2.0* prediction results with MetaSRA.** The boxplots show the log2(auPRC/prior) scores for the overall (**a**) and breakdown (**b**) of tissue classification performance for *txt2onto 2.0* (word + LR) and MetaSRA, as well as combined performance. Only non-redundant tasks that are shared between *txt2onto 2.0* and MetaSRA are used to assess the performance. Each point in the boxplots corresponds to the performance of a single tissue model. The grey dashed line represents the median performance of the *txt2onto 2.0* model. In breakdown performance, tasks are divided into two quantiles with an equal number of tasks based on the number of positive instances in the training set.

(a)

(b)

(c)

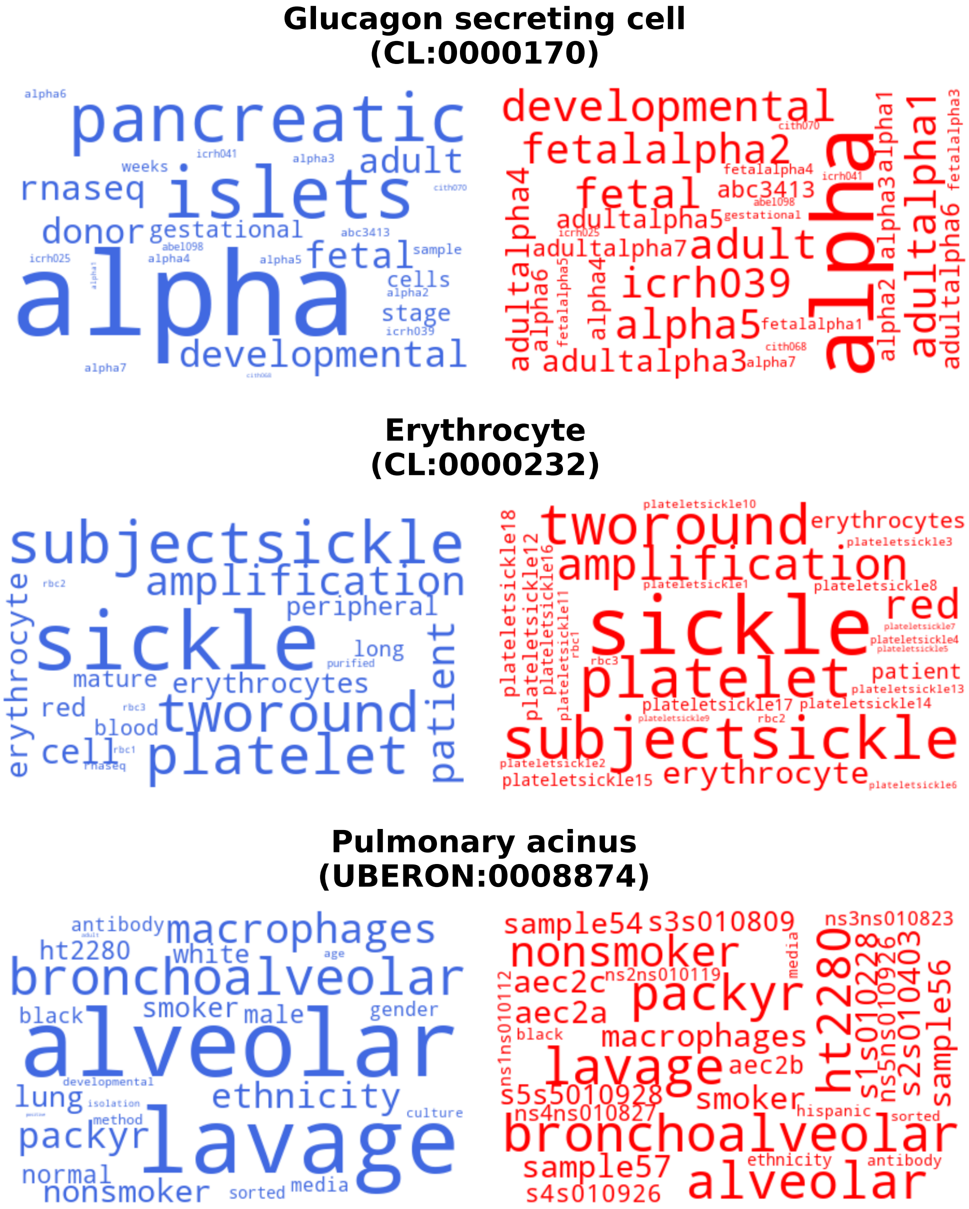


Figure S7: **Predictive words extracted from LR and PM models for tissue classification**. The word clouds show the predictive words extracted from three top-performing tissue classification models. The words in royal blue are from the LR models, while those in red are from the corresponding PM models. In the word clouds generated from the LR model, the size of each word is proportional to its regression coefficient. In the word clouds generated from the PM model, the size of each word is proportional to its performance score ($\log_2$(auPRC/prior)) in predicting labels.

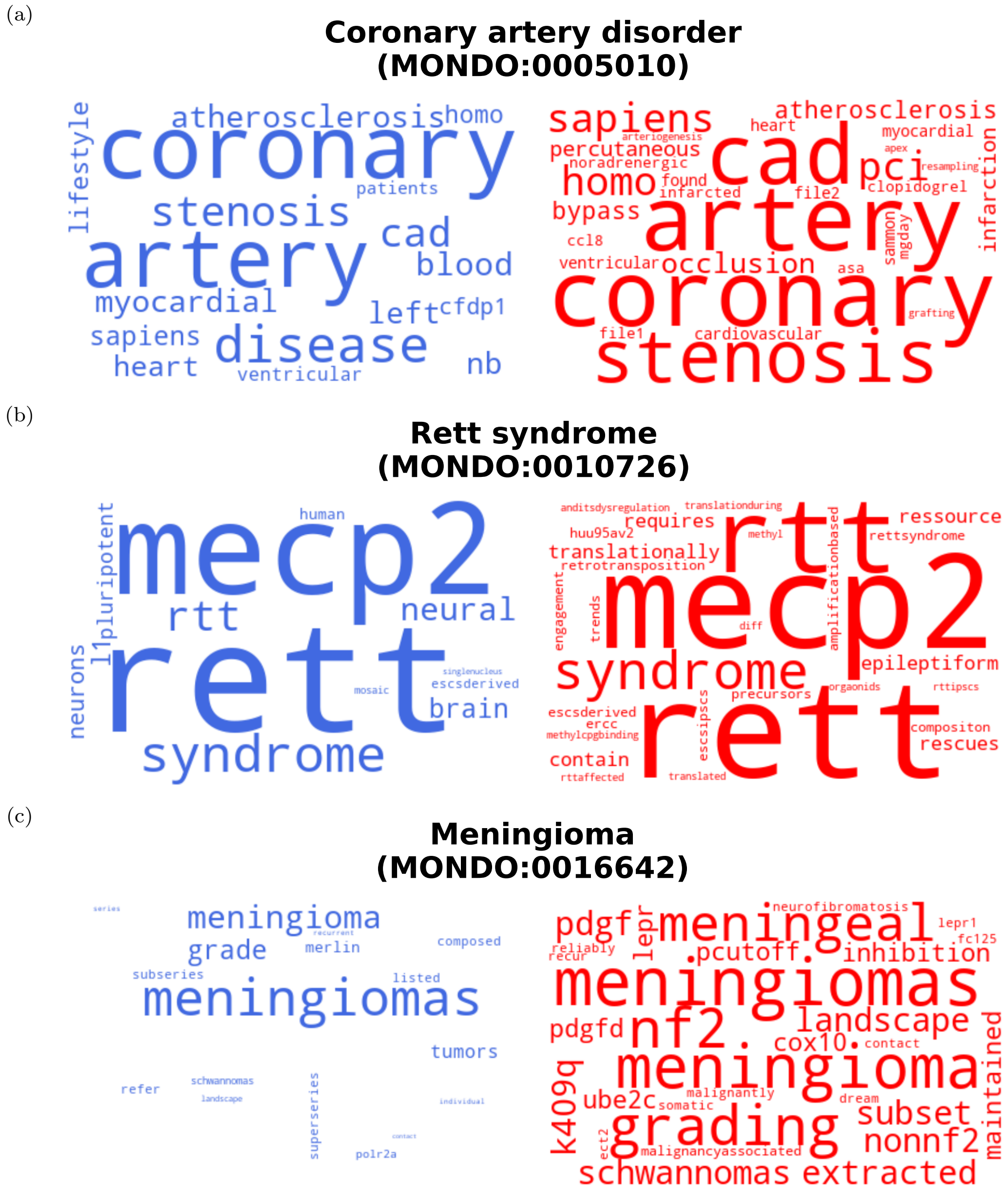


Figure S8: **Predictive words extracted from LR and PM models for disease classification**. The word clouds show the predictive words extracted from three top-performing disease classification models. The words in royal blue are from the LR models, while those in red are from the corresponding PM models. In the word clouds generated from the LR model, the size of each word is proportional to its regression coefficient. In the word clouds generated from the PM model, the size of each word is proportional to its classification performance score ($\log_2$(auPRC/prior)) in predicting labels.

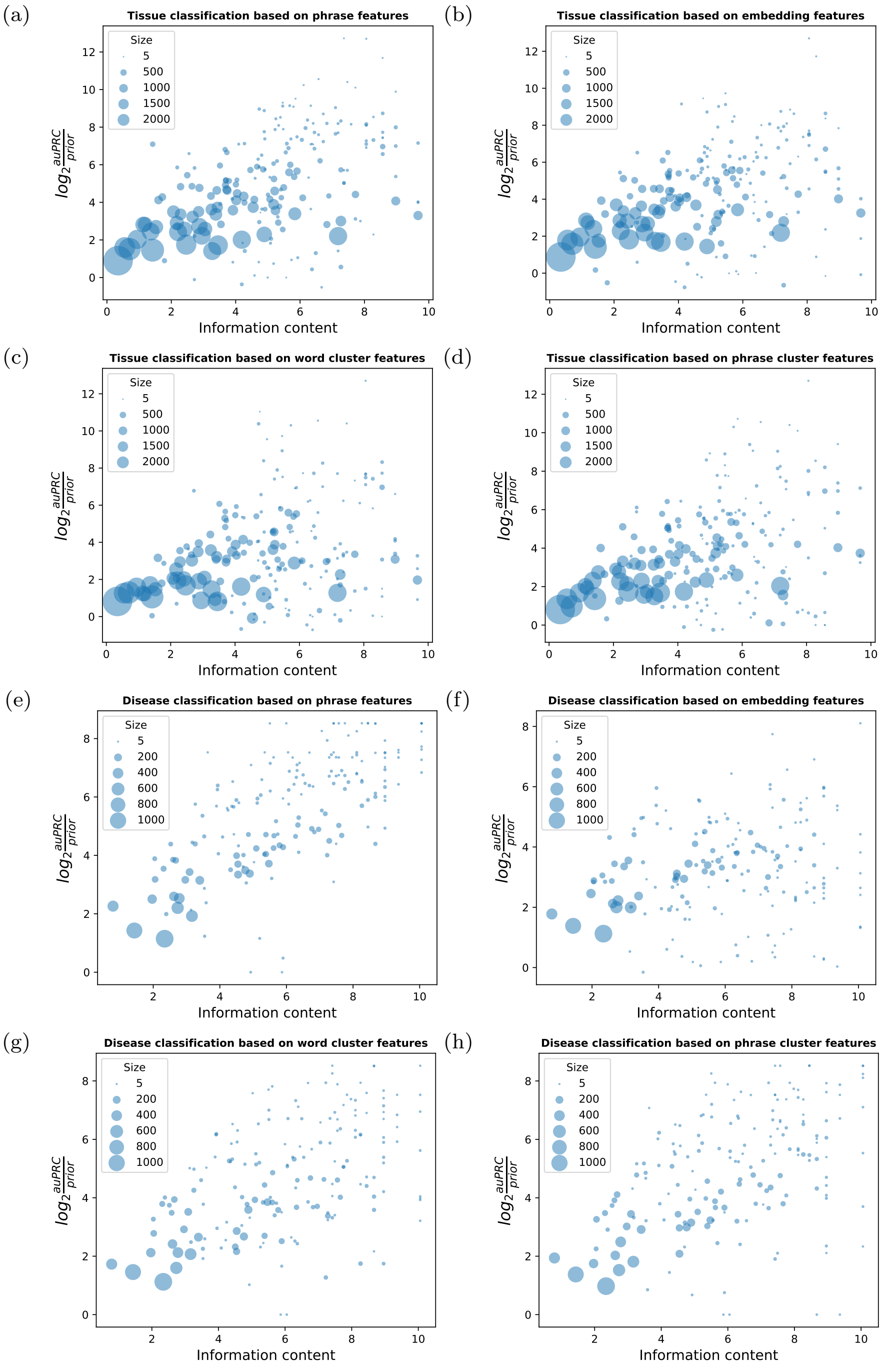


Figure S9: **Effect of term specificity on prediction performance across models build on various features**. Scatter plots show the prediction performance of the logistic regression model for disease (**e, f, g, h**) and tissue (**a, b, c, d**) classification tasks across various levels of term specificity. Models are trained based on various features including phrase (**a, e**), word embedding (**b, f**), word cluster (**c, g**) and phrase cluster (**d, h**). Each point represents the performance of a single tissue or disease model. The x-axis shows the specificity of the disease or tissue terms, defined by their information content (IC). The IC of a term is the negative logarithm of the fraction of terms in the ontology that are descendants of term of interest. The y-axis shows the model performance, measured by log2(auPRC/prior). The size of each data point is proportional to the number of positive instances in the training set for each task.

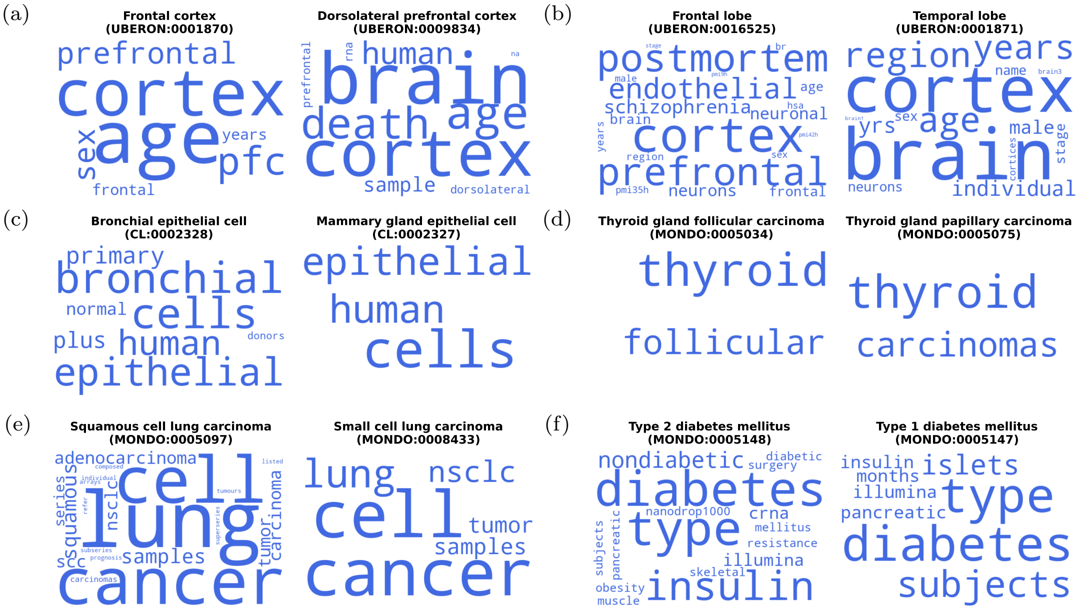


Figure S10: **Semantically similar terms that are challenging to differentiate**. The word clouds present pairs of tissue (**a, b, c**) or disease terms (**d, e, f**) that are semantically similar, but cannot be differentiated by the best-performing model (logistic regression model with word features). The word clouds in each panel comprise task-related words extracted from the metadata of testing instances annotated to these terms. The size of each word in the clouds is proportional to its frequency among the task-related words, with a larger size indicating a higher prevalence.

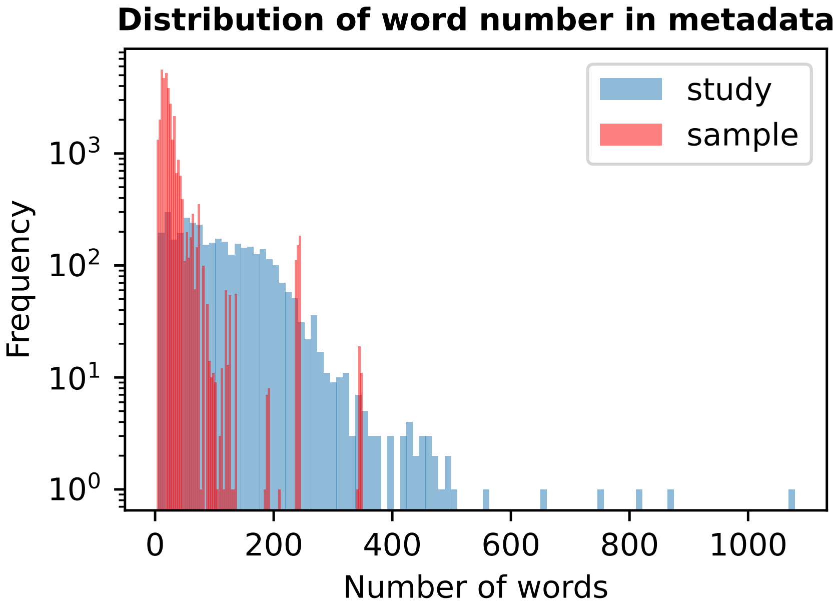


Figure S11: **Distribution of word number in metadata**. The histograms show the distribution of word numbers in sample-level (red) and study-level (blue) metadata. The frequency on the y-axis is displayed on a logarithmic scale.

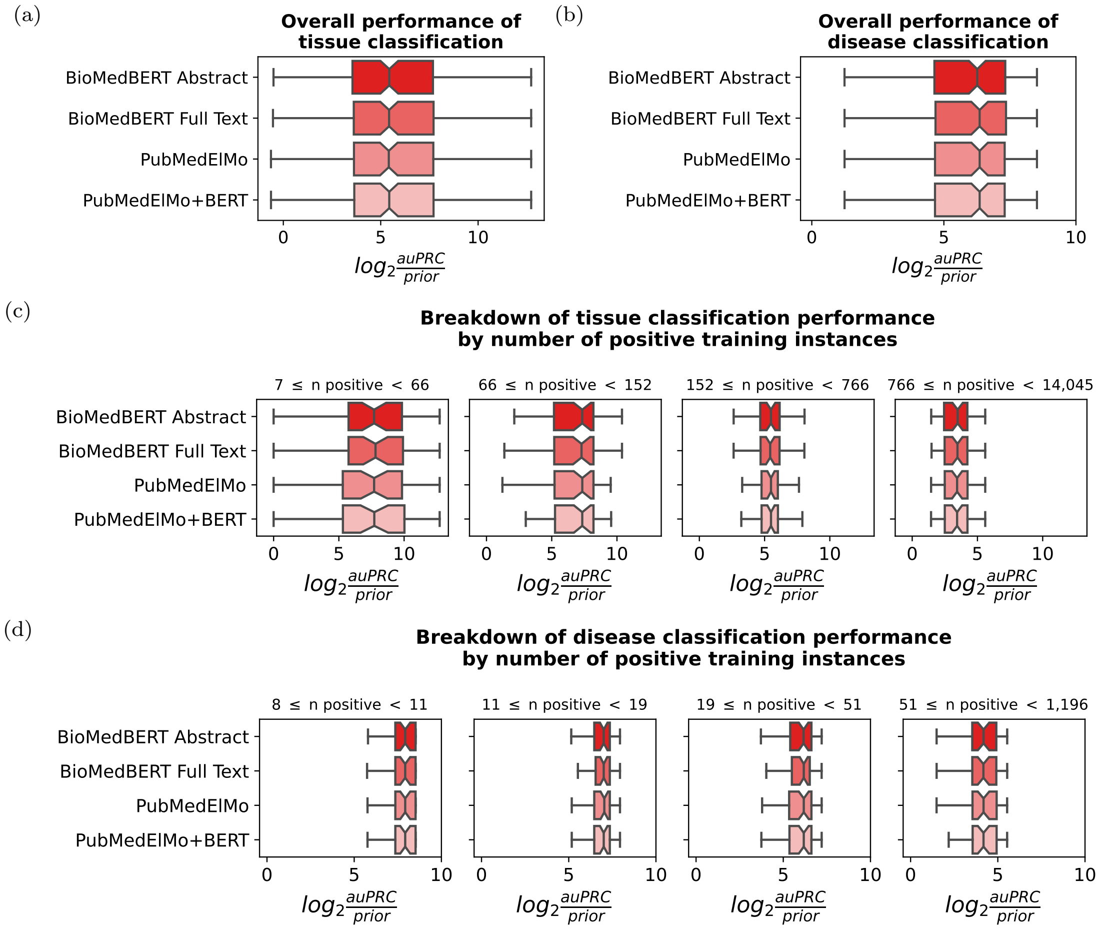


Figure S12: **Effect of word embeddings on model performance.** The boxplots show the effect of applying word embeddings generated from different language models on the tissue and disease classification performance, measured by $\log_2$(auPRC/prior). The predictions are made by the best performing model (logistic regression model with word features), while using different embeddings as input, including BioMedBERT pretained on abstract (BioMedBERT Abstract), BioMedBERT pretained on full text (BioMedBERT Full Text), PubMed ElMo and stack embedding of PubMedElMo and BERT model pretained on generic text (PubMedElMo + BERT). (**a, b**) The boxplots show the effect of word embeddings on overall performance of tissue (a) and disease (b) classification. (**c, d**) The boxplots show the breakdown of tissue (c) and disease (d) classification performance. Tasks are divided into four quantiles with an equal number of tasks based on the number of positive instances in the training set.

#### Supplementary Tables

| | ***Txt2onto 1.0*** | ***Txt2onto 2.0*** |
|---|---|---|
| Feature vectors | Embedding vector of the entire description | TF-IDF vector of the description |
| Model explainability | No | Yes |
| Handling unseen text during prediction | Embedding entire description into the same latent space | Using the word embedding space to map unseen tokens to semantically similar seen tokens |
| Tissue annotation | Yes | Yes |
| Disease annotation | No | Yes |
| Num of trained tissue models | 346 | 590 |
| Num of trained disease models | NA | 1,166 |

Table S1: **Model comparison between *txt2onto 1.0* and *2.0*.**

| **Source** | **Num of samples** | **Num of tissue types** |
|---|---|---|
| Hawkins et al. | 12,218 | 81 |
| In-house | 5,025 | 11 |
| TissueNexus | 6,296 | 15 |
| Sirota et al. | 1,530 | 41 |
| Gu et al. | 5,677 | 93 |
| Bernstein et al. | 3,103 | 293 |
| **Total** | 33,849 | 440 |

Table S2: **Summary of datasets used for tissue classification models.** This table shows the number of studies and disease types from datasets used for training and evaluating disease classification models. The total in the table represents the total number of study and disease types after removing shared samples and types among the sources. As a result, the numbers from each source do not add up to the total.

| **Source** | **Num of studies** | **Num of disease types** |
|---|---|---|
| In-house | 421 | 47 |
| USRA-HD | 99 | 253 |
| Sirota et al. | 67 | 56 |
| Gu et al. | 123 | 116 |
| GEMMA | 2,966 | 902 |
| **Total** | 3,676 | 1,096 |

Table S3: **Summary of datasets used for disease classification models.** This table shows the number of samples and tissue types from datasets used for training and evaluating tissue classification models. The total in the table represents the total number of samples and tissue types after removing shared samples and types among the sources. As a result, the numbers from each source do not add up to the total.

| ClinicalTrials ID | MONDO ID | Name | P | log2(P/prior) | Recal P |
|---|---|---|---|---|---|
| NCT04024644 | MONDO:0005076 | periodontitis | 0.0047 | 1.79 | 1.00 |
| NCT02385474 | MONDO:0005076 | periodontitis | 0.0041 | 1.59 | 1.00 |
| NCT04680806 | MONDO:0005076 | periodontitis | 0.0038 | 1.50 | 1.00 |
| NCT03448107 | MONDO:0005076 | periodontitis | 0.0038 | 1.47 | 1.00 |
| NCT03176537 | MONDO:0005076 | periodontitis | 0.0037 | 1.45 | 1.00 |
| NCT06051396 | MONDO:0005149 | pulmonary hypertension | 0.0158 | 3.54 | 1.00 |
| NCT03269630 | MONDO:0005149 | pulmonary hypertension | 0.0098 | 2.85 | 1.00 |
| NCT00125918 | MONDO:0005149 | pulmonary hypertension | 0.0084 | 2.62 | 1.00 |
| NCT02587325 | MONDO:0005149 | pulmonary hypertension | 0.0079 | 2.53 | 1.00 |
| NCT00825266 | MONDO:0005149 | pulmonary hypertension | 0.0079 | 2.53 | 1.00 |
| NCT06095791 | MONDO:0004981 | atrial fibrillation | 0.0125 | 3.20 | 1.00 |
| NCT05928728 | MONDO:0004981 | atrial fibrillation | 0.0120 | 3.14 | 1.00 |
| NCT01970969 | MONDO:0004981 | atrial fibrillation | 0.0116 | 3.09 | 1.00 |
| NCT00142194 | MONDO:0004981 | atrial fibrillation | 0.0115 | 3.08 | 1.00 |
| NCT04496336 | MONDO:0004981 | atrial fibrillation | 0.0111 | 3.03 | 1.00 |
| NCT00115622 | MONDO:0011786 | allergic rhinitis | 0.0057 | 2.07 | 1.00 |
| NCT00894231 | MONDO:0011786 | allergic rhinitis | 0.0051 | 1.90 | 1.00 |
| NCT00691665 | MONDO:0011786 | allergic rhinitis | 0.0050 | 1.87 | 1.00 |
| NCT03431961 | MONDO:0011786 | allergic rhinitis | 0.0046 | 1.75 | 1.00 |
| NCT01279057 | MONDO:0011786 | allergic rhinitis | 0.0044 | 1.70 | 1.00 |
| NCT02538809 | MONDO:0004986 | urinary bladder carcinoma | 0.0159 | 3.54 | 1.00 |
| NCT03832803 | MONDO:0004986 | urinary bladder carcinoma | 0.0135 | 3.30 | 1.00 |
| NCT04720222 | MONDO:0004986 | urinary bladder carcinoma | 0.0096 | 2.81 | 1.00 |
| NCT03463915 | MONDO:0004986 | urinary bladder carcinoma | 0.0095 | 2.79 | 1.00 |
| NCT00491114 | MONDO:0004986 | urinary bladder carcinoma | 0.0089 | 2.71 | 1.00 |
| NCT04849559 | MONDO:0005036 | gastric adenocarcinoma | 0.0165 | 3.58 | 1.00 |
| NCT03484949 | MONDO:0005036 | gastric adenocarcinoma | 0.0154 | 3.49 | 1.00 |
| NCT02541461 | MONDO:0005036 | gastric adenocarcinoma | 0.0146 | 3.41 | 1.00 |
| NCT06042998 | MONDO:0005036 | gastric adenocarcinoma | 0.0145 | 3.40 | 1.00 |
| NCT02466711 | MONDO:0005036 | gastric adenocarcinoma | 0.0135 | 3.30 | 1.00 |

Table S4: **Top predictions for diseases from Clinicaltrial studies.** This table shows the top five predictions from the top six best-performing disease classification models. Each row corresponds to a study. The first column is the MONDO ID for the disease term of the classification model, the second column is the corresponding disease name, the third column is the uncalibrated probability, the fourth column is the uncalibrated probability normalized by the prior and then log2 transformed, and the final column is the recalibrated probability.

| PRIDE ID | MONDO ID | Name | P | log2(P/prior) | Recal P |
|---|---|---|---|---|---|
| PXD015931 | MONDO:0005132 | cytomegalovirus infection | 0.0052 | 1.92 | 1.00 |
| PXD005276 | MONDO:0005132 | cytomegalovirus infection | 0.0040 | 1.57 | 1.00 |
| PXD023559 | MONDO:0005132 | cytomegalovirus infection | 0.0040 | 1.55 | 1.00 |
| PXD009945 | MONDO:0005132 | cytomegalovirus infection | 0.0038 | 1.46 | 1.00 |
| PXD013120 | MONDO:0005132 | cytomegalovirus infection | 0.0038 | 1.46 | 1.00 |
| PXD013988 | MONDO:0006486 | uveal melanoma | 0.0064 | 2.19 | 1.00 |
| PXD023511 | MONDO:0006486 | uveal melanoma | 0.0051 | 1.87 | 1.00 |
| PXD029132 | MONDO:0006486 | uveal melanoma | 0.0050 | 1.82 | 1.00 |
| PXD011883 | MONDO:0006486 | uveal melanoma | 0.0046 | 1.72 | 1.00 |
| PXD014368 | MONDO:0006486 | uveal melanoma | 0.0044 | 1.64 | 1.00 |
| PXD025629 | MONDO:0100096 | COVID-19 | 0.0732 | 4.75 | 1.00 |
| PXD023450 | MONDO:0100096 | COVID-19 | 0.0610 | 4.48 | 1.00 |
| PXD018581 | MONDO:0100096 | COVID-19 | 0.0587 | 4.43 | 1.00 |
| PXD022789 | MONDO:0100096 | COVID-19 | 0.0556 | 4.35 | 1.00 |
| PXD041281 | MONDO:0100096 | COVID-19 | 0.0531 | 4.28 | 1.00 |
| PXD012173 | MONDO:0005184 | pancreatic ductal adenocarcinoma | 0.1606 | 4.87 | 1.00 |
| PXD015492 | MONDO:0005184 | pancreatic ductal adenocarcinoma | 0.1516 | 4.79 | 1.00 |
| PXD030879 | MONDO:0005184 | pancreatic ductal adenocarcinoma | 0.1242 | 4.50 | 1.00 |
| PXD031910 | MONDO:0005184 | pancreatic ductal adenocarcinoma | 0.1063 | 4.28 | 1.00 |
| PXD032951 | MONDO:0005184 | pancreatic ductal adenocarcinoma | 0.1046 | 4.25 | 1.00 |
| PXD009241 | MONDO:0010150 | head and neck squamous cell carcinoma | 0.2041 | 5.22 | 1.00 |
| PXD001438 | MONDO:0010150 | head and neck squamous cell carcinoma | 0.1348 | 4.62 | 1.00 |
| PXD001862 | MONDO:0010150 | head and neck squamous cell carcinoma | 0.0912 | 4.06 | 1.00 |
| PXD007705 | MONDO:0010150 | head and neck squamous cell carcinoma | 0.0707 | 3.69 | 1.00 |
| PXD030343 | MONDO:0010150 | head and neck squamous cell carcinoma | 0.0594 | 3.44 | 1.00 |
| PXD021201 | MONDO:0004953 | invasive ductal breast carcinoma | 0.1828 | 5.00 | 1.00 |
| PXD027012 | MONDO:0004953 | invasive ductal breast carcinoma | 0.1024 | 4.16 | 1.00 |
| PXD037900 | MONDO:0004953 | invasive ductal breast carcinoma | 0.0945 | 4.05 | 1.00 |
| PXD029020 | MONDO:0004953 | invasive ductal breast carcinoma | 0.0910 | 3.99 | 1.00 |
| PXD037920 | MONDO:0004953 | invasive ductal breast carcinoma | 0.0891 | 3.96 | 1.00 |

Table S5: **Top predictions for diseases from PRIDE studies.** This table shows the top five predictions from the top six best-performing disease classification models. Each row corresponds to a study. The first column is the MONDO ID for the disease term of the classification model, the second column is the corresponding disease name, the third column is the uncalibrated probability, the fourth column is the uncalibrated probability normalized by the prior and then log2 transformed, and the final column is the recalibrated probability.

### Supplementary Methods

#### Preparing gold standard

##### Collecting labeled examples and constructing gold standard

We obtained tissue labels for samples from GEMMA [1], Hawkins *et al.* [2], Gu *et al.* [3], Bernstein *et al.* [4], Sirota *et al.* [5], TissueNexus [6] and an in-house curated dataset, covering 33,849 samples and 440 tissue types. If samples overlapped among these datasets, labels were prioritized in the following order: Hawkins *et al.* > in-house > TissueNexus > Sirota *et al.* > Gu *et al.* > Bernstein *et al.*. We obtained disease labels for studies from multiple sources, including GEMMA (accessed on 7/9/2023) [1], Gu *et al.* [3], Sirota *et al.* [5], USRA-HD [7], and an in-house curated dataset, covering 3,676 studies and 1,069 disease types. If there were duplicated studies included, we prioritized the annotations in the following order: in-house > USRA-HD > Sirota *et al.* > Gu *et al.* > GEMMA. All tissue and disease annotations were mapped to MONDO and UBERON-CL, respectively.

Next, we used the annotations of samples and studies to build a gold standard following the procedure described in Lee *et al.* [8]. For every term, if a sample or study was directly annotated to the term or was a descendant of the term, it was labeled as positive. If a sample or study was annotated to an ancestor of a term, it was labeled as unknown since the annotation was ambiguous. The rest of the samples or studies were labeled as negative. Samples or studies from GEMMA and Bernstein *et al.* might be labeled to multiple terms. For those cases, if a sample or study could be labeled as positive based on at least one of the terms, it was labeled as positive. For samples or studies that could not be labeled as positive, they were labeled as negative if at least one of the terms was labeled as negative. The remaining cases were labeled as unknown.

##### Selecting terms for model evaluation

As described above, since we propagated labels along the ontology, positive labels of closely related parent-child terms might largely overlap. These redundant tasks could potentially bias the performance score distribution by inflating the scores for overlapping terms. To mitigate this issue, we found groups of redundant terms by comparing shared positive instances and then selecting representative terms from each group. The procedure is conducted as follows. First, we measured the similarity of positive instance sets between terms using the Jaccard index. Then, we constructed a network of terms in which two terms were connected if their Jaccard index exceeded the threshold, meaning the pair of terms shared a significant number of positive instances. We set 0.9 and 0.5 as the minimum Jaccard index threshold for tissue and disease, respectively. Next, we extracted all connected components from the term network, and each connected component contained redundant terms. To extract a set of representative terms from the connected components, we first ordered the terms in descending order based on the number of neighbors they have. For each term, we formed a cluster with the term and its neighbors. For each cluster, we selected a representative term that was annotated to the highest number of instances in the original sample or disease labels before propagation, then removed the entire cluster from the connected component. We repeated this process until all connected components were depleted. Notably, connected components could contain only one term, so we did not miss terms unrelated to any other terms.

To guarantee robust training and evaluation, we selected a subset of non-redundant terms with sufficient positive instances for training. For disease terms, we first split the dataset into training and testing sets in an 80/20 ratio. The training set was further split into at least two folds for hyperparameter tuning. To ensure unbiased model evaluation, the difference in the positive-to-negative ratio between the training and testing sets was controlled to be under 0.1. This same criterion was applied to the ratio difference between folds. The data splitting for tissue terms also required all the above criteria. Additionally, samples from the same study must be kept within the same set or fold to avoid data leakage.

Among all disease and tissue/cell terms in the ontology, 590 tissue and 1,166 disease terms were trainable. After removing redundant tasks, 237 non-redundant tissue/cell terms and 353 disease terms remained suitable for training. These tasks can be used for training but may not have enough positive instances for hyperparameter tuning. Of these trainable terms, 202 tissue/cell and 177 disease terms had sufficient instances in the training sets to be further split into at least two folds, enabling hyperparameter tuning through cross-validation.

#### Downloading metadata from GEO

GEO houses comprehensive study and sample descriptions, so we aimed to download all metadata from GEO [9]. Since samples from Gu *et al.* and Bernstein *et al.* were named with Sequence Read Archive (SRA) IDs, we used the efetch tool version 17.0 [10] to map SRA sample IDs to GEO sample IDs. Samples not included in GEO were removed from the analysis. Similarly, we used the efetch tool to obtain the GEO study ID for each sample. Then, we downloaded the soft format file for each study directly from NCBI and extracted all text related to the sample and study descriptions.

#### *Txt2onto 2.0*

##### Preprocessing input

Preprocessing has a crucial effect on the model's ability to make accurate predictions. In addition to basic preprocessing steps such as removing non-UTF-8 characters, URLs, punctuation, whole numbers, and converting text to lowercase, we tested several additional preprocessing settings that might impact prediction performance. One preprocessing step involved removing task-irrelevant fields, as information from those fields might confound model training and prediction. For tissue classification, we treated "Sample_title", "Sample_source_name_ch1", "Sample_description" and "Sample_characteristics_ch1" as task-relevant fields. For disease classification, we treated "Series_overall_design", "Series_summary", and "Series_title" as task-relevant fields. We found that removing task-irrelevant fields does affect model performance (**Fig. S1**). Lemmatization is another widely applied technique for preprocessing text, which aims to reduce the feature space by converting words to their original root form. Lemmatization has a limited effect on model performance (**Fig. S2**). However, ineffective lemmatization could yield meaningless strings, especially for biomedical words. Hence, we opted to keep the text in its original form. Additionally, we defined a pool of text snippets to be removed from the corpus, primarily consisting of names of biomolecular reagent companies and reagent kits.

##### Converting preprocessed text into TF-IDF matrix

After preprocessing, we converted the preprocessed text into a TF-IDF matrix. TF-IDF is a popular method for extracting information from free text, measuring how frequently a word is observed in the document (term frequency, TF) and how unique this term is across the document corpus (inverse document frequency, IDF). Since the training and testing sets may not share the same corpus, the TF-IDF matrix features are calculated separately for each set.

For the training set, the TF-IDF was calculated as follows:

$$\text{TF-IDF}(t, d, D) = \text{TF}(t, d) \times \log \frac{D}{df_t} \quad (1)$$

For a given metadata in the training set, $\text{TF}(t, d)$ was the term frequency of a word $t$ within that metadata $d$. $D$ was the total number of documents in the training set, and $df_t$ was the frequency of this word $t$ occurring across the entire training corpus $D$. For each feature in the testing set, we aligned it to the most similar feature in the training set by comparing its cosine similarity of text embedding to training features. If multiple testing features were mapped to the same training feature, their term frequencies were summed up, and the inverse document frequency was inherited from corresponding features in the training set. After normalizing the TF-IDF by instance, we obtained TF-IDF matrix of a testing set with the same features as the training set. The similarity between training and testing features was measured as the cosine similarity between their embeddings. We tested the effect of selecting different language models and found that the choice of embedding has a limited impact on performance (**Fig. S12**). We selected the base uncased version of BioMedBERT [11] pretrained on abstracts to generate embeddings using PyTorch 2.1.0 [12] and the Transformer 4.36.2 Python library [13].

Since individual words may not fully capture the meaning of biomedical terms, we also tried using phrases as features. In this case, TF was the phrase frequency, and IDF was the uniqueness of this phrase across the corpus. To reduce redundant entities, we built a K-nearest neighbor graph, where each node was an entity, and edges were the cosine similarity between entities ($K = 1$). Clusters were then found using Leiden community detection [14]. The TF of a cluster was calculated as the sum of TF of all entities in the cluster, and the IDF was the inverse frequency of at least one entity in the cluster being observed across the corpus.

##### Training the LR classifiers

We chose the logistic regression model as the predictive model due to its superior performance compared to other classifiers in *txt2onto 1.0* [2], and its output probability is well-calibrated. ElasticNet was selected as the penalty term since it can reduce the influence of features unrelated to the prediction while keeping features with similar meanings [15]. This is particularly useful when dealing with sparse matrices like the TF-IDF matrix. The model was trained using scikit-learn version 1.2.1 [16]. We conducted cross-fold validation training to determine the best hyperparameters. Based on the results of hyperparameter tuning (see **Supplementary file 1**), we built the model with the following parameters: $C = 1$, $alpha = 0.4$, $penalty = elasticnet$, and $solver = liblinear$. Notably, despite the model yielding the best performance when $C = 1.0$ and $alpha = 0.0$ in most cases, we raised alpha to 0.4, since we wanted the model to penalize features to zero while maintaining comparable performance when $C = 1.0$ and $alpha = 0.0$. By setting alpha to 0.4, the model can effectively regularize features to zero while maintaining a performance level similar to the optimal settings.

##### Evaluating classification results

We measured classification performance on the testing set using $\log_2$(auPRC/prior), the auPRC normalized by the prior, which is the ratio of positive instances to the total instances in the training set. This normalization addresses class imbalance, given that positive samples are often much fewer than negative ones for most tissue or disease terms. Normalization also allows for consistent comparison across different tissue and disease terms, despite variations in the distribution of positive samples.

##### Differentiating between similar terms

We compared the model's performance when differentiating similar and dissimilar terms. Semantic similarity between terms was measured by the cosine similarity between the text embeddings of each pair of terms' names, which was generated by BioMedBERT. For every pair of terms, we used the trained model from one of the terms to predict the probability of instances annotated to these two terms in the testing set. We treated the instances annotated to the same terms as the task of the model as positive instances, then treated the instances related to the other term in the pair as negative instances. We expected the probability of the positive instances to be higher than the negative instances, and measured the performance of differentiating positive and negative instances by auROC. We did not use $\log_2$(auPRC/prior) as in assessing the general performance of the models since the ratio between positive and negative instances is not proportional to the one in the training set, so the prior cannot inform random classification performance in this case. Additionally, since labels of instances have been propagated to general terms, an instance could be labeled to multiple terms. We removed instances that appeared in both positive and negative sets. We also did not include low-performance models with $\log_2$(auPRC/prior) lower than two in the analysis, as they cannot effectively distinguish among terms. Finally, we included 408 terms and 92,517 pairs of terms for disease classification, and 237 terms and 39,361 pairs of terms for tissue classification.

##### Predicting on independent datasets

Besides annotating metadata in GEO, we also wanted to test the performance of our model when predicting on independent datasets. We chose ClinicalTrials and PRIDE as test cases. We downloaded metadata for all studies from ClinicalTrials (accessed on 12/6/2023) and PRIDE (accessed on 1/22/2024). For efficient testing, we randomly selected 20,000 and 10,000 studies from ClinicalTrials and PRIDE, respectively, for disease annotation. Data were preprocessed in the same way as described for GEO metadata. We employed the LR model using words as features for prediction, which achieved the best performance over the rest of the entity types. Since the ratio of positive and negative instances was highly imbalanced in some tasks, the output probability from the model might be underestimated. We trained another Logistic Regression model using the probability before calibration and ground truth label from the testing set without penalty terms. Then, we applied it to recalibrate the prediction probability from independent datasets. Recalibrated probabilities above 0.5 were considered positive predictions. Another useful metric for indicating good predictions was $log_2(probability/prior)$, which measured how much better the predicted probability was compared to the probability of random predictions. If this score exceeded 1, the prediction was considered a trustworthy positive prediction.

We also conducted a double-blind test to evaluate model performance on independent datasets. For each dataset, we selected six top-performing models with at least twenty positive predictions and at least three positive instances during training. Potential positive predictions were selected based on $log_2(probability/prior)$, considering instances as potential positives if the value exceeded 1. Then, we selected the top and bottom ten predictions and ordered them according to $log_2(probability/prior)$ in descending order, where the top ten were expected to be positive, and the bottom ten predictions were expected to be negative. We divided these predictions into two sets, each containing five positive and negative predictions. For the first set, everything remained the same, while for the second set, we kept the study order but shuffled the labels, purposely making some random predictions. Finally, we combined the two sets and shuffled the order of the studies with their labels. By comparing the performance of predictions from the first set, where the study and predicted label matched, to the second set, which represented random predictions, we can quantitatively evaluate the performance of predictions on independent datasets. The shuffled list was sent to three experts for manual evaluation. The averaged performance scores were used as the final score.

#### Combining *txt2onto 2.0* and MetaSRA predictions

We used precomputed MetaSRA predictions that were downloaded from the following link: `https://metasra.biostat.wisc.edu/static/metasra_versions/v1.8/metasra.v1-8.json`. Sample IDs from the MetaSRA predictions (SRA accession codes) were mapped to GSMs using mappings from the SRA archive obtained here: `ftp://ftp.ncbi.nlm.nih.gov/sra/reports/Metadata`. Next, we combined the mapped MetaSRA predictions with our gold-standard tissue labels, leaving 14,248 overlapping sample-level predictions.

Predictions from MetaSRA and *txt2onto 2.0* were combined via a weighted average of each prediction using model performance from a training set as weights. We used auPRC as the performance metric. We required that each training set had at least ten positive examples and at least three for each test set, leaving 68 tissue and cell type terms for evaluation. We did not evaluate predictions for disease tasks since there were not enough positive examples shared between MetaSRA predictions and our curated labels.

### Supplementary Note 1

In this section, we provide a step-by-step description for *txt2onto 2.0*.

1. Preprocess text
   1.1 Remove non-UTF-8 characters from the text
   1.2 Remove URLs from the document content
   1.3 Remove punctuation marks and symbols
   1.4 Remove numerical values
   1.5 Convert all remaining text to lowercase format
   1.6 Remove fields not relevant to the analysis task
   1.7 Combine text from remaining fields into a single document
2. Partition dataset
   2.1 Split dataset into training and testing set by 80/20 ratio
   2.2 For tissue classification, keep samples from the same study in the same partition
3. Engineer training set feature
   3.1 Create training vocabulary:
      - Extract all unique words from training documents, which are treated as training feature set
   3.2 Calculate feature values:
      - Calculate term frequencies (TF) of every feature in each document
      - Calculate inverse document frequency (IDF) of every feature in each document
      - Combine TF and IDF to get TF-IDF scores
4. Engineer testing set feature
   4.1 Align features:
      - Extract all unique words from testing documents, which are testing feature set
      - Calculate cosine similarity between word embeddings of every pair of training and testing feature
      - Identify most similar training feature for each testing feature
   4.2 Generate TF-IDF matrix:
      - Calculate TF of testing feature in each document
      - Map TF of testing features to corresponding training features based on cosine similarity
      - Sum TF when multiple testing features map to same training feature
      - Calculate TF-IDF using mapped TF and training feature IDF
5. Training model
   5.1 Use training set TF-IDF matrix as input features
   5.2 Use corresponding labels as target values
   5.3 Train ElasticNet model
6. Evaluate model
   6.1 Input testing set TF-IDF matrix into trained model to calculate prediction probabilities for each document
   6.2 Calculate performance score based on corresponding labels

#### Supplementary Note 2

In addition to the logistic regression model (LR), we proposed a baseline model called the power matrix (PM) to investigate how word or other entity features are predictive of labels. PM measures the overall relevance of a given text to the term. The intuition behind PM is that if a feature's TF-IDF value is higher in positive instances than in negative ones, it is potentially related to the given term. This is equivalent to assessing the performance of using the TF-IDF value of a feature to predict labels. We used the training set exclusively to evaluate the features' predictive performance. We measured this performance using $\log_2$(auPRC/prior), where auPRC is normalized by *prior*. *Prior* is the proportion of positive instances in the training set, representing the expected performance of a random classifier. This metric reflects how much better a feature can predict labels compared to random classification within the training set. To measure the overall relevance of a given text in the testing set, we calculated the score of each instance as the weighted average of the performance scores, weighted by the TF-IDF of the corresponding features. Similar to regression coefficients in LR model, the performance measure of text features in PM is also highly interpretable, where higher value indicate better predictive power of the feature.

Comparing LR with PM, we find that the performance of both models is generally on par (**Fig. S4**). In scenarios with a limited number of positive samples for both tissue and disease tasks, PM tends to slightly outperform LR (**Fig. S5**). This difference in performance may be attributed to PM's higher sensitivity in capturing weak signals from predictive features, which might be regularized to zero in LR models.

We also wanted to determine whether PM can capture biologically meaningful entities. Compared to other entities, the PM model with word features demonstrates the best performance; thus, we focused solely on predictive word features. We considered all words with positive $\log_2$(auPRC/prior) values as predictive word features. By comparing predictive words from PM and LR, we find that predictive words from LR are generally a subset of words from PM (**Fig. S7,S8**). Careful examination of extra predictive words generated by PM reveals that PM tends to capture uninformative words with small $\log_2$(auPRC/prior) values. For example, "sample56" in *Pulmonary acinus* (UBERON:0008874) (**Fig. S7c**) and "file2" in *Coronary artery disorder* (MONDO:000510) (**Fig. S8a**) are random words unrelated to the terms. The PM also tends to capture more correlated words than LR. For instance, PM captures the word "plateletsickle" ending with various numbers in the model for *Erythrocyte* (CL:0000232) (**Fig. S7b**). These correlated words also have small $\log_2$(auPRC/prior) values. Capturing uninformative or too many correlated words is potentially due to the fact that, unlike the LR model, the PM model does not have any regularization on uninformative and highly correlated features. However, regularization is a double-edged sword. Without regularization, the PM sometimes captures more informative words than LR. For example, the PM model for *Pulmonary acinus* (UBERON:0008874) highlights "aec2a", "aec2b" and "aec2c", which represent a cell type, alveolar secretory cell, within the *Pulmonary acinus* (**Fig. S7c**). The PM model for *Meningioma* (MONDO:0016642) captures "pdgf", representing platelet-derived growth factor, which is expressed in meningioma patients (**Fig. S8c**).

Despite the excellent performance of the PM model and its high sensitivity to task-related words, we still chose LR over PM since regularization in LR can exclude uninformative features, resulting in a more biologically interpretable model, which is the primary goal of *txt2onto 2.0*. Another reason is that the output of PM is the weighted average of performance scores, not a probability, making it less intuitive to evaluate the quality of the outcome compared to LR.

Nevertheless, the strong performance of the PM model as a baseline for metadata classification confirms that accurate disease and tissue predictions can be effectively achieved by identifying predictive words from the metadata, rather than requiring a comprehensive understanding of the entire text. This finding highlights the potential of simple, interpretable models in biomedical text classification tasks, particularly when dealing with limited positive samples.

### Supplementary Note 3

In this section, we provide examples of specific disease and tissue terms that exhibited poor performance in our analysis. We also present the study IDs of the positive instances for disease terms and the study IDs from which the positive samples originated for tissue terms. Furthermore, we discuss potential reasons for the poor prediction performance observed for these terms.

1. Example of disease terms with poor classification performance
   - *AIDS* (MONDO:0012268):
     - Positive training instances: GSE56484, GSE17372
     - Positive testing instances: GSE17491
     - Reason: The predictive words in the training set - “hiv” and “hiv1” - do not exist in the testing set. The word indicating AIDS in the positive testing instance is “hivinfected”, however, the closest word to “hivinfected” in the training features is neither “hiv” nor “hiv1”.
   - *Acute promyelocytic leukemia* (MONDO:0012883):
     - Positive training instances: GSE12662, GSE2550
     - Positive testing instances: GSE5949
     - Reason: The positive instance in the testing set is general leukemia, but not acute promyelocytic leukemia.
   - *Gliosarcoma* (MONDO:0016681):
     - Positive training instances: GSE166696, GSE29796
     - Positive testing instances: GSE8692
     - Reason: Positive training instances are studies about glioma, not gliosarcoma.
   - *Stroke disorder* (MONDO:0005098):
     - Positive training instances: GSE158314, GSE161450
     - Positive testing instances: GSE180722
     - Reason: The positive testing instance is a study related to alcohol use disorder, not stroke.
2. Example of tissue terms with poor classification performance
   - *Hematopoietic stem cell* (CL:0000037):
     - Positive training instances: GSE19429, GSE29524, GSE19680, GSE2049, GSE76234, GSE75384, GSE55689, GSE19240, GSE29523
     - Positive testing instances: GSE60
     - Reason: Positive testing instances in this study were sourced from lymphoma and thus were mislabeled as hematopoietic stem cells.
   - *Proximal-distal subdivision of colon* (UBERON:0000168):
     - Positive training instances: GSE46513, GSE1710
     - Positive testing instances: GSE65107
     - Reason: Positive training instances were sourced from the colon, but testing instances were coming from colorectal tissue, which encompasses both the colon and the rectum.
   - *Retina* (UBERON:0000966):
     - Positive training instances: GSE22765, GSE40524
     - Positive testing instances: GSE2705
     - Reason: Positive testing instances were sourced from lamina cribrosa cells, which is a structure within the optic nerve head of the eye. They were mislabeled as the retina.

- *Exocrine gland of integumental system* (UBERON:0019319):
  - Positive training instances: GSE83083, GSE10046, GSE18070, GSE73628, GSE12790
  - Positive testing instances: GSE5060
  - Reason: Positive testing instances were sourced from airway epithelium, which is mislabeled as nipple (UBERON:0002030).

### Supplementary Note 4

Below, we list several examples of similar disease pairs that are challenging for *txt2onto 2.0* to distinguish. We also provide studies related to these terms. For disease terms, we include the IDs of relevant studies. Additionally, we discuss potential reasons why *txt2onto 2.0* struggles to differentiate between these disease pairs.

1. Examples of similar disease pairs that cannot be teased apart by *txt2onto 2.0*

   - *Type 2 diabetes mellitus* (MONDO:0005148) vs. *Type 1 diabetes mellitus* (MONDO:0005147)
     - Positive instances: GSE40234, GSE29231, GSE21340, GSE27951
     - Negative instances: GSE214851, GSE60424, GSE24147, GSE33440
     - AUC: 0.56
     - Cosine similarity: 0.99
     - Reason: Whole numbers were removed during preprocessing, which make model difficult to distinguish between Type 1 and Type 2 diabetes.
   - *Squamous cell lung carcinoma* (MONDO:0005097) vs. *Small cell lung carcinoma* (MONDO:0008433)
     - Positive instances: GSE42998, GSE28582, GSE20853, GSE7880, GSE43580, GSE21933
     - Negative instances: GSE31625, GSE164247
     - AUC: 0.58
     - Cosine similarity: 0.99
     - Reason: GSE28582 and GSE21933 should be labeled to non-small cell lung carcinoma, a more general disease than squamous cell lung carcinoma. GSE20853 should be labelled as another subtype of non-small cell lung carcinoma - lung adenocarcinoma.
   - *Follicular lymphoma* (MONDO:0018906) vs. *Mantle cell lymphoma* (MONDO:0018876)
     - Positive instances: GSE118707, GSE37088
     - Negative instances: GSE10793
     - AUC: 0.50
     - Cosine similarity: 0.99
     - Reason: GSE118707 should be labelled as another type of lymphoma, Hodgkins' lymphoma.
   - *Thyroid gland follicular carcinoma* (MONDO:0005034) vs. *Thyroid gland papillary carcinoma* (MONDO:0005075)
     - Positive instances: GSE32662,GSE61844
     - Negative instances: GSE29265,GSE83520
     - AUC: 0.25
     - Cosine similarity: 0.99
     - Reason: The thyroid gland papillary carcinoma studies, GSE29265 and GSE83520, are generally predicted to have a higher probability of being classified as thyroid gland follicular carcinoma compared to the actual thyroid gland follicular carcinoma studies, GSE32662 and GSE61844. This discrepancy in classification may be attributed to the shared term ”thyroid gland” in both papillary and follicular carcinoma descriptions.
